# Genetic Grouping of SARS-CoV-2 Coronavirus Sequences using Informative Subtype Markers for Pandemic Spread Visualization

**DOI:** 10.1101/2020.04.07.030759

**Authors:** Zhengqiao Zhao, Bahrad A. Sokhansanj, Charvi Malhotra, Kitty Zheng, Gail L. Rosen

## Abstract

We propose an efficient framework for genetic subtyping of SARS-CoV-2, the novel coronavirus that causes the COVID-19 pandemic. Efficient viral subtyping enables visualization and modeling of the geographic distribution and temporal dynamics of disease spread. Subtyping thereby advances the development of effective containment strategies and, potentially, therapeutic and vaccine strategies. However, identifying viral subtypes in real-time is challenging: SARS-CoV-2 is a novel virus, and the pandemic is rapidly expanding. Viral subtypes may be difficult to detect due to rapid evolution; founder effects are more significant than selection pressure; and the clustering threshold for subtyping is not standardized. We propose to identify mutational signatures of available SARS-CoV-2 sequences using a population-based approach: an entropy measure followed by frequency analysis. These signatures, Informative Subtype Markers (ISMs), define a compact set of nucleotide sites that characterize the most variable (and thus most informative) positions in the viral genomes sequenced from different individuals. Through ISM compression, we find that certain distant nucleotide variants covary, including non-coding and ORF1ab sites covarying with the D614G spike protein mutation which has become increasingly prevalent as the pandemic has spread.

ISMs are also useful for downstream analyses, such as spatiotemporal visualization of viral dynamics. By analyzing sequence data available in the GISAID database, we validate the utility of ISM-based subtyping by comparing spatiotemporal analyses using ISMs to epidemiological studies of viral transmission in Asia, Europe, and the United States. In addition, we show the relationship of ISMs to phylogenetic reconstructions of SARS-CoV-2 evolution, and therefore, ISMs can play an important complementary role to phylogenetic tree-based analysis, such as is done in the Nextstrain [1] project. The developed pipeline dynamically generates ISMs for newly added SARS-CoV-2 sequences and updates the visualization of pandemic spatiotemporal dynamics, and is available on Github at https://github.com/EESI/ISM and via an interactive website at https://covid19-ism.coe.drexel.edu/.

**Author Summary:** The novel coronavirus responsible for COVID-19, SARS-CoV-2, expanded to reportedly 8.7 million confirmed cases worldwide by June 21, 2020. The global SARS-CoV-2 pandemic highlights the importance of tracking viral transmission dynamics in real-time. Through June 2020, researchers have obtained genetic sequences of SARS-CoV-2 from over 47,000 samples from infected individuals worldwide. Since the virus readily mutates, each sequence of an infected individual contains useful information linked to the individual’s exposure location and sample date. But, there are over 30,000 bases in the full SARS-CoV-2 genome—so tracking genetic variants on a whole-sequence basis becomes unwieldy. We describe a method to instead efficiently identify and label genetic variants, or “subtypes” of SARS-CoV-2. Applying this method results in a compact, 11 base-long compressed label, called an Informative Subtype Marker or “ISM”. We define viral subtypes for each ISM, and show how regional distribution of subtypes track the progress of the pandemic. Major findings include (1) covarying nucleotides with the spike protein which has spread rapidly and (2) tracking emergence of a local subtype across the United States connected to Asia and distinct from the outbreak in New York, which is found to be connected to Europe.

## Introduction

Severe acute respiratory syndrome coronavirus 2 (SARS-CoV-2), the novel coronavirus responsible for the COVID-19 pandemic, was first reported in Wuhan, China in December 2019. [2, 3]. In a matter of weeks, SARS-CoV-2 infections had been detected in nearly every country, and as of July 2020, reported cases continue to rapidly increase across multiple continents. Powered by advances in rapid genetic sequencing, there is an expansive and growing body of data on SARS-CoV-2 sequences from individuals around the world. During the early stage of the pandemic, a substantial degree of heterogeneity was already identified, with differences in 15% of the sites of the sequences. [4] SARS-CoV-2 will mutate over time as transmissions occur and the virus spreads; although, notably, it has previously been observed that coronaviruses, which are single strand RNA viruses with a relatively large genome size ("30,000 bases), tend to have lower mutation rates than other RNA viruses [5]. Central repositories are continuously accumulating SARS-CoV-2 genome data from around the world, such as the Global Initiative on Sharing all Individual Data (GISAID) [6] (available at https://www.gisaid.org/).

Researchers are presently using whole genome sequence alignment and phylogenetic tree construction to study the evolution of SARS-CoV-2 on a macro and micro scale [1, 7–10]. For example, the Nextstrain group has created a massive phylogenetic tree incorporating sequence data and applied a model of the time-based rate of mutation to create a hypothetical map of viral distribution [1] (available at https://nextstrain.org/ncov). Similarly, the China National Center for Bioinformation has established a “2019 Novel Coronavirus Resource”, which includes a clickable world map that links to a listing of sequences along with similarity scores based on alignment (available at https://bigd.big.ac.cn/ncov?lang=en) [11].

In more granular studies, early work by researchers based in China, analyzing 103 genome sequences, identified two highly linked single nucleotides, leading them to suggest that two major subtypes had emerged: one called “L,” predominantly found in the Wuhan area, and “S,” which derived from “L” and found elsewhere [12]. Subsequently, further diversity was recognized as the virus continued to spread, and researchers developed a consensus reference sequence for SARS-CoV-2, to which other sequences may be compared [13]. Researchers have continued to publish studies of the specific variants in the context of localized outbreaks, such as the *Diamond Princess* cruise ship [14], as well as regional outbreaks and their international connections [12, 15–18].

Efforts are also underway to identify potential genome sites and regions where selection pressure may result in phenotypic variation. Particular focus has been given to the ORF (open reading frame) coding for the spike (S) receptor-binding protein, which may impact the development of vaccines and antivirals [19]. Notably, a group studying sequence variants within patients reported limited evidence of intrahost variation, though they cautioned that the results were preliminary and could be the result of limited data [20, 21]. Intrahost variation thus represents yet another layer of complexity in evaluating that viral variation which influences disease progression in an individual patient, or may be associated with events that can in turn generate sequence variation in other individuals that patient infects.

Given the importance of tracking and modeling genetic changes in the SARS-CoV-2 virus as the outbreak expands, there is a need for an efficient methodology to quantitatively characterize groups of variation in the SARS-CoV-2 virus genome by defining genetic subtypes of the virus. Exemplary potential applications of quantitative subtyping include the following: 1) Characterizing potentially emerging variants of the virus in different regions, which may ultimately express different phenotypes. 2) Monitoring variation in the viral genome that may be important for vaccine development, for example due to emerging structural differences in proteins encoded by different strains. 3) Designing future testing methodology to contain disease transmission across countries and regions, for example developing specific tests that can characterize whether a COVID-19 patient developed symptoms due to importation or likely domestic community transmission. 4) Identifying viral subtypes that may correlate with different clinical outcomes in different regions and patient subpopulations.

Phylogenetic trees obtained through sequence alignment may be utilized to map viral outbreaks geographically and trace transmission chains [22, 23] and have been applied to SARS-CoV-2 by, e.g., the Nextstrain group as discussed above. At an early stage in the pandemic, however, phylogenetic trees may be unreliable predictors of evolutionary relationships between viral strains circulating worldwide, because of insufficient information regarding the molecular clock assumption, practical limits on data collection, and sampling bias [24]. Accordingly, subtyping based on phylogenetic models may also be unreliable and change, as the assumptions underlying the models change given more sequencing and continued variation in the viral genome. The ISM approach described in this paper relies instead on compact measures of sequence similarity that will remain conserved even as more genome sequence data is added over time. ISMs thus provide a robust subtype definition, which can help track the virus as the pandemic progresses. Therefore, it may be more efficient to focus on co-occurring patterns of only the sites of the more frequently occurring variation within the viral genome to identify subtypes, rather than utilizing whole genome sequence data to cluster viral genomes, which may contain additional confounding variation.

Moreover, the nomenclature of clades imply that the categorization of viruses in subtypes is static rather than dynamic. SARS-CoV-2 is a novel virus in humans that is rapidly evolving, which makes it harder to establish a stable nomenclature for genetic typing [10]. The Nextstrain project has sought to address these challenges by providing their own clade definitions based on whether there are a certain number of mutations at nucleotide positions in the sequence (at least two) and naming clades based on their estimated time of emergence [25]. This shows that a conventional genetic subtype relying on whole genome phylogenetic trees will be complicated by the changes in viral genome, especially early on in a pandemic before those changes are clearly governed by selection pressure. Lineages will likely disappear and reemerge within different geographical regions and over the course of time [10]. In addition, viral evolutionary analysis, such as by the Nextstrain group, relies on making assumptions solely on molecular evolution (the degree of sequence similarity and branching points in defining genetic clades and other levels of organization) and not on transmission models.

In this paper, we propose a methodology to complement phylogenetics-based transmission and evolution models of SARS-CoV-2 that can consistently and rapidly identify subtypes without requiring an initial tree reconstruction step—and, thereby, avoid the need to make assumptions about the molecular evolution clock and clustering thresholds. To generate highly informative molecular signatures indicative of a subtype or emerging lineage, we look to methods that have been successfully employed in the microbiome field to resolve species/subspecies from 16S ribosomal RNA (16S rRNA) gene [26]. The 16S rRNA gene is a highly conserved sequence and therefore can be used for phylogenetic analysis in microbial communities [27–31]. One way to differentiate between closely related microbial taxa is to identify nucleotide positions in 16S rRNA data (“oligotypes”) that represent information-rich variation [32]. This approach has also been used in the reverse direction to find conserved sites as a way to assemble viral phylogenies [33]. Dawy et al. proposed to use Shannon’s mutual information to identify multiple important loci for Gene mapping and marker clustering [34]. Shannon Entropy [35] has been applied in multiple sequence alignment data to quantify the sequence variation at different positions [32, 36]. Given a position of interest, entropy can be used to measure the amount of “randomness” at that position, as determined by whether sequences may have different bases at a specific position. For instance, if there is an A at a given position across all aligned sequences, the entropy will be 0, i.e., there is no “randomness” at that position. On the other hand, if at a given position there is a G in 50% of the sequences and a T in the other 50%, the entropy will be 1 (i.e., essentially “random”), and thus a relatively high entropy. Based on this property, oligotyping [32] utilizes variable sites revealed by the entropy analysis to identify highly refined taxonomic units.^1^

Accordingly, we present herein a method to define a genetic signature, called an “informative subtype marker” or ISM, for the viral genome that can be 1) utilized to define SARS-CoV-2 subtypes that can be quantified to characterize the geographic and temporal spread of the virus, and 2) efficiently implemented for identifying strains to potentially analyze for phenotypic differences. The method compresses the full viral genome to generate a small number of nucleotides that are highly informative of the way in which the viral genome dynamically changes. We draw on the aforementioned oligotyping approach developed for 16S rRNA data [32] and build on its implementation of entropy and grouping patterns to address the particular challenges of viral genomes. On top of oligotyping, we add error correction to account for ambiguities in reported sequence data and, optionally, applied further compression by identifying patterns of base entropy correlation. The resulting ISM, therefore, defines a viral genetic subtype (that can be related to a phylogenetic “lineage”, see Comparison of ISM-defined subtypes to clades identified using phylogenetic trees) in the sense that it is a compressed (reduced complexity) representation of a set of genetic features (a.k.a genotype).

The ISM pipeline may complement a phylogenetic approach in that it can efficiently identify viral subtypes of the population through genetic hotspots and do not rely on evolutionary model assumptions. ISMs identify subtypes with slight differences between sequences where the sequence identity is >99%, as is the case of SARS-CoV-2 with OrthoANI of 99.8% (at the end of April 2020) [37]. ISMs include the key base mutations in the marker identification itself. And thus, unlike phylogenetic lineages (i.e., clades and subsequent emergent subtypes), ISM-defined subtypes are expressly differentiated by mutations with high diversity (over the viral population). For example, the ISM label of a subtype can include mutation in SARS-CoV-2’s spike protein, which may have an important phenotypic impact.

As a succinct and robust identifier, therefore, ISM-based subtyping can facilitate downstream analysis, such as modeling and visualizing the geographic and temporal patterns of genetic variability of SARS-CoV-2 sequences obtained from the GISAID database. We have made the pipeline available on Github https://github.com/EESI/ISM, where it will be continuously updated as new sequences are uploaded to data repositories ^2^. We have also developed an interactive website showing the worldwide country-specific distributions of ISM-defined subtypes, available at https://covid19-ism.coe.drexel.edu/

## Methods

### Data collection and preprocessing

Nextstrain maintains a continually-updated, pre-formatted SARS-CoV-2 (novel coronavirus) sequence dataset through GISAID (this dataset also includes sequences of other novel coronavirus sampled from other hosts such as Bat). This dataset was downloaded from GISAID (http://www.gisaid.org) on June 17, 2020, which contains 47,305 sequences. Our preprocessing pipeline then begins by filtering out sequences that are less than 25000 base pairs (the same threshold used in Nextstrain project built for SARS-CoV-2^3^). We also included a reference sequence from National Center for Biotechnology Information^4^ (NCBI Accession number: NC 045512.2), resulting in an overall data set of 47,280 sequences. We then performed multiple sequence alignment on all remaining sequences using MAFFT [38] with the “FFT-NS-2” method in XSEDE [39]. After alignment, the sequence length was extended (for the present data set, up to 79716 nt) due to gaps inserted into the sequences during the multiple sequence alignment.

### Entropy analysis and ISM extraction

For the aligned sequences, we merged the sequence with the metadata provided by Nextstrain^5^ as of June 17, 2020, based on identification number, **gisaid epi isl**, provided by GISAID [6]. Given the fast-moving nature of the pandemic, we filtered out sequences with incomplete date information in metadata (e.g. “2020-01”) in order to incorporate temporal information with daily resolution. In addition, we filtered out sequences from unknown host or non-human hosts. The resulting final data set contained 45535 sequences excluding the reference sequence. Then, we calculated the entropy at a given position *i* by:

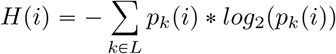

where *L* is a list of unique characters in all sequences and *p*_*k*_(*i*) is a probability of observing a character *k* at position *i*. We estimated *p*_*k*_(*i*) from the frequency of characters at that position. We refer to characters in the preceding because, in addition to the bases `A, C, G`, and T, the sequences include additional characters representing gaps (-) and ambiguities, which are listed in Supplementary file 2 — Sequence notation [40].^6^

Bases like `N` and -, which represent a fully ambiguous site and a gap respectively, are substantially less informative. Therefore, we further define a *masked entropy* as entropy calculated without considering sequences containing `N` and - in a given nucleotide position in the genome. With the help of this masked entropy calculation, we can focus on truly informative positions, instead of positions at the start and end of the sequence in which there is substantial uncertainty due to artifacts in the sequencing process. Finally, high entropy positions are selected by two criteria: 1) entropy > 0.23, and 2) the percentage of `N` and - is less than 25%. Further details about the selection of these two criteria are provided in Supplementary file 1 — Masked entropy threshold analysis. In the data set we processed for this paper, the entropy threshold yielded 20 distinct positions within the viral genome sequence. We built the Informative Subtype Markers (ISMs) at these 20 nucleotide positions on each sequence.

### Error correction to resolve ambiguities in sequence data and remove spurious ISMs

The focus of the error correction method is to resolve an ISM that contains ambiguous symbols, i.e., a nucleotide identifier that represents an ambiguous base call (as detailed in Supplementary file 2 — Sequence notation [40]), such as N, which represents a position that could be `A, C, T`, or `G`. Our approach uses ISMs with few or no ambiguous symbols to correct ISMs with many ambiguities. Given an ISM with an error, we first find all ISMs that are identical to the subject ISM’s nucleotide positions without error. We refer here to these nearly-identical ISMs as supporting ISMs. Then, we iterate over all positions with an error that must be corrected in the subject ISM. For a given nucleotide position, if all other such supporting ISMs with respect to the said erroneous position contain the same *non-ambiguous* base (i.e., an `A, C, T`, or `G)`, then we simply correct the ambiguous base to that *non-ambiguous* base found in the supporting ISMs. However, when the supporting ISMs disagree at a respective nucleotide position, the method generates an ambiguous symbol which represents all the bases that occurred in the supporting ISMs and compare this artificially generated nucleotide symbol with the original position in the subject ISM. If the generated nucleotide symbol identifies a smaller set of bases, e.g., `Y` representing `C` or `T` rather than `N`, which may be any base, then we use the generated symbol to correct the original one.

When we applied the foregoing error correction algorithm to ISMs generated from the genome data set analyzed in this paper, we found that 90.2% of erroneous ISMs were partially corrected (meaning at least one nucleotide position with ambiguity was corrected for that ISM if not all), and 24.5% of erroneous ISMs were fully corrected (meaning all positions with ambiguity were corrected to a *non-ambiguous* base (i.e., an `A, C, T`, or `G`)). Since one ISM may represent multiple sequences in the data set, overall the error correction algorithm was able to partially correct 96.0% of sequences identified by an erroneous ISM, and 32.4% of such sequences were fully corrected.

The error correction method necessarily results in the replacement of ISMs with an ambiguous base at a site by another ISM without an error at that site. We expect, and have observed that the abundance of non-ambiguous ISMs are inflated by the error correction process. Here, we utilize the inflation rate of ISMs to quantify the difference in abundance of an ISM before and after error correction process. The inflation rate is defined by:

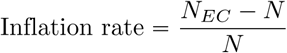

where *N* is the abundance of an ISM of interest (typically an ISM with few or no ambiguous bases) before error correction, and *N*_*EC*_ is the abundance of that ISM after error correction.

### Quantification and visualization of viral subtypes

At the country/region level, we assess the geographic distribution of SARS-CoV-2 subtypes, and, in turn, we count the frequency of unique ISMs per location and build charts and tables to visualize the ISMs, including the pie charts, graphs, and tables shown in this paper. All visualizations in this paper and our pipeline are generated using Matplotlib and Plotly [41, 42]. To improve visualization, ISMs that occur with frequency of less than 5% in a given location are collapsed into “OTHER” category per location. Our pipeline then creates pie charts for different locations to show the geographical distribution of subtypes. Each subtype is also labeled with the earliest date associated with sequences from a given location in the dataset.

The abundance table based ordination is widely used to visualize community ecology in the microbiome [43]. [44] also used Principal Components Analysis (PCA) to produce a two-dimensional visual summary of the genetic variation in human populations. In our application, we can use the abundances of different ISMs in a country as features to quantify the genetic variation pattern of SARS-CoV-2 sequences. In our analysis, we select countries that have more than 100 viral sequences uploaded in order to have enough ISMs to viably generate such an abundance table. Then, the number of sequences is down-sampled to 100 for each country/region with more than 100 sequences so that all countries/regions have the same effective “sequencing depth.” Therefore, results are not biased by the different number of submissions in different countries. We then construct the ISM abundance table. The elements in the abundance table represent the abundance of an ISM in a country/region after down-sampling, where each column is an ISM. We use Bray-Curtis dissimilarity [45] to quantify the dissimilarity of ISM compositions between a pair of regions and form a pairwise Bray-Curtis dissimilarity matrix. Finally, we employ PCA to reduce the dimensionality of the pairwise Bray-Curtis dissimilarity matrix, plotting the first two components to visualize the genetic variation patterns of those countries/regions.

To study the progression of SARS-CoV-2 viral subtypes in the time domain, we group all sequences in a given location that were obtained no later than a certain date (as provided in the sequence metadata) together and compute the relative abundance (i.e., frequency) of corresponding subtypes. Any subtypes with a relative abundance that never goes above 2.5% for any date are collapsed into “OTHER” category per location. The following formula illustrates this calculation:

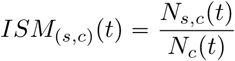

where *ISM*_(*s,c*)_(*t*) is the relative abundance of a subtype, *s*, in location, *c*, at a date *t, N*_*s,c*_(*t*) is the total number of instances of such subtype, *s*, in location, *c*, that has been sequenced no later than date *t* and *N*_*c*_(*t*) is the total number of sequences in location, *c*, that has been sequenced no later than date *t*.

### Comparison of ISM subtyping to phylogenetic analysis

Nextstrain [1] provides a phylogeny method to track and visualize the dynamic of SARS-CoV-2 sequences. To obtain the results presented here, we downloaded the Nextstrain tree data from https://nextstrain.org/ncov on June 17, 2020. Since both the ISMs and the Nextstrain phylogenetic tree were generated based on the GISAID database, they may be easily compared. We present two comparisons in this paper: 1) ISM hamming distance and phylogenetic tree branch length; 2) ISM clusters defined at different entropy thresholds and Nextstrain defined “clades”.

The highest-abundance ISMs are involved with hundreds, if not thousands, of sequences. For a given ISM, we can find the lowest common ancestor (LCA) node in the phylogenetic tree of all sequences with that ISM. The branch length between the LCA and the root can be considered as the inferred evolutionary distance between the reference sequence and the LCA node. Hamming distance between our ISMs measures the divergence between two clusters of sequences. We can compare branch length between the root and LCA with the Hamming distance between a given ISM and the reference ISM. Then, we compute the Pearson correlation coefficient to measure the correlation between the evolutionary distance from the reference genome (inferred by the phylogenetic tree) and the Hamming distance between ISMs and the reference ISM.

In Nextstrain data, there are 5 “clades”, namely, 19A, 19B, 20A, 20B and 20C, defined in [25]. Different sequences are assigned to those 5 “clades” based on genetic variations in the sequence. Since our ISMs are clusters of sequences with similar genetic variations, the Nextstrain “clades” provides us a good interface to study how our entropy threshold influences the ISM definition by comparing the overlaps between Nextstrain “clades” grouped sequences and ISM grouped sequences. To measure the similarity between ISM labels and “clade” labels of sequences, we use two clustering metrics, homogeneity and completeness as proposed in [46]. A clustering result satisfies homogeneity if all of its clusters contain only data points which are members of a single class. A clustering result satisfies completeness if all the data points that are members of a given class are elements of the same cluster [46]. We vary the entropy threshold to form different sets of ISM clusters of sequences and compare each set with Nextstrain “clades” using homogeneity and completeness.

## Results and Discussion

We begin by identifying and mapping the sites that form an ISM for each genome based on sequence entropy. Then, we analyze the properties of ISMs and validate the ISMs generated from SARS-CoV-2 data as of June 17, 2020. We present ISM abundance inflation introduced by error correction, demonstrate how ISMs evolve as a function of entropy threshold, and show how entropy values at different positions change over time. Then, we show the visualization of spatiotemporal dynamics based on ISMs. We analyze the geographic distribution of SARS-CoV-2 genetic subtypes identified by ISMs, as well as the temporal dynamics of the subtypes. We also visualize the viral genetic variation patterns of different regions based on their ISM subtype abundances. Then, we evaluate the results of ISM subtyping in comparison with current genetic variation studies of SARS-CoV-2. Finally, we compare the ISM subtypes to viral “clades” that were determined by Nextstrain, in order to demonstrate how ISMs relate to evolutionary relationships predicted by phylogenetic methods.

### Identification and Mapping of Subtype Markers

In this section, we briefly discuss the potential functional relevance of the identified ISM locations. We further demonstrate that minimal artifacts are introduced by the error correction methodology, which indicates that ISM identification is stable with respect to the choice of entropy threshold within a reasonable range. Finally, we generate compressed ISM labels based on correlated entropy variation between ISM sites.

#### Identification of ISM locations by whole genome sequence entropy analysis

The first step in the ISM subtyping pipeline is the determination of the entropy at each nucleotide position in the SARS-CoV-2 genome in order to identify the sites that will make up the ISM. Entropy is used to quantify the variation at different positions for sequence alignment result. For a given position with high entropy value, there are more than 1 nucleotides showing up frequently at this position across all the aligned sequences. On the other hand, if the entropy value is low at a position, it implies that this position is more conserved across all aligned sequences. Figure 1 shows the overall entropy at each nucleotide position, determined based on calculating the masked entropy for all sequences as described in the Methods section. Notably, at the beginning and end of the sequence, there is a high level of uncertainty. This is because there are more N and - symbols, representing ambiguity and gaps, in these two regions (gaps are likely a result of artifacts in MAFFT’s alignment of the viruses or its genomic rearrangement [21], and both ambiguous base-calls (N’s) and gaps (-’s) may result due to the difficulty of accurately sequencing the genome at the ends). After applying filtering to remove low entropy positions and uncertain positions, we identified 20 informative nucleotide positions in the sequence to generate informative subtype markers (see filtering details in Methods section).

**Figure 1.**
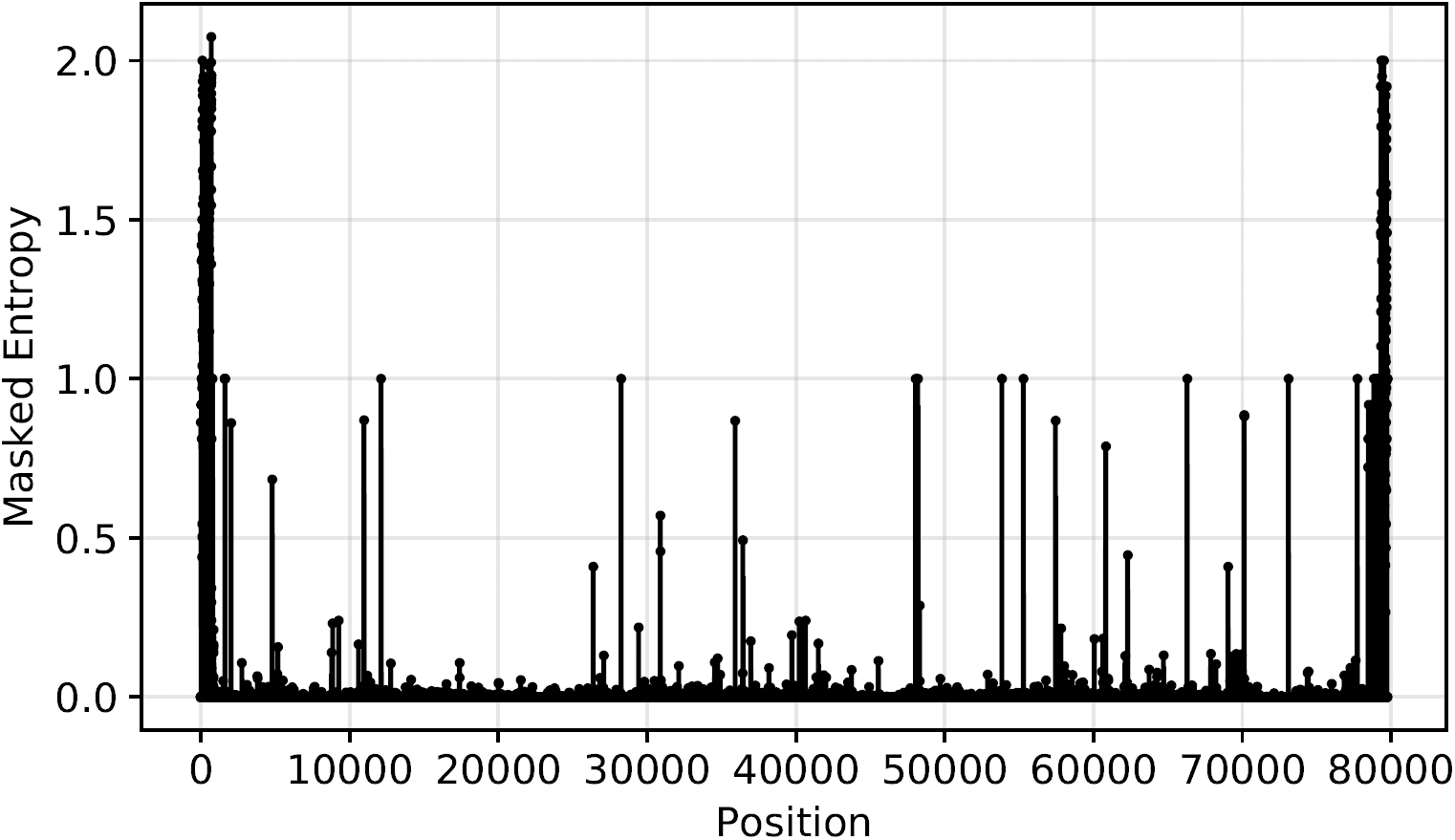
Overall entropy as a function of nucleotide position for all SARS-CoV-2 sequences in the data set. The peaks in this figure corresponds to highly variable positions and positions with 0 or lower entropy values represent conservative regions in the aligned viral genomes.

Importantly, even though the combinatorial space for ISM is potentially very large due to the substantial number of characters that may present at any one nucleotide position, only certain ISMs occur in substantial quantities in the overall sequence population. Figure 2 demonstrates the rapid decay of the frequency of sequences with a given ISM. In particular, the plot shows that the first three ISMs represent subtypes that have more than 4000 sequences worldwide.

**Figure 2.**
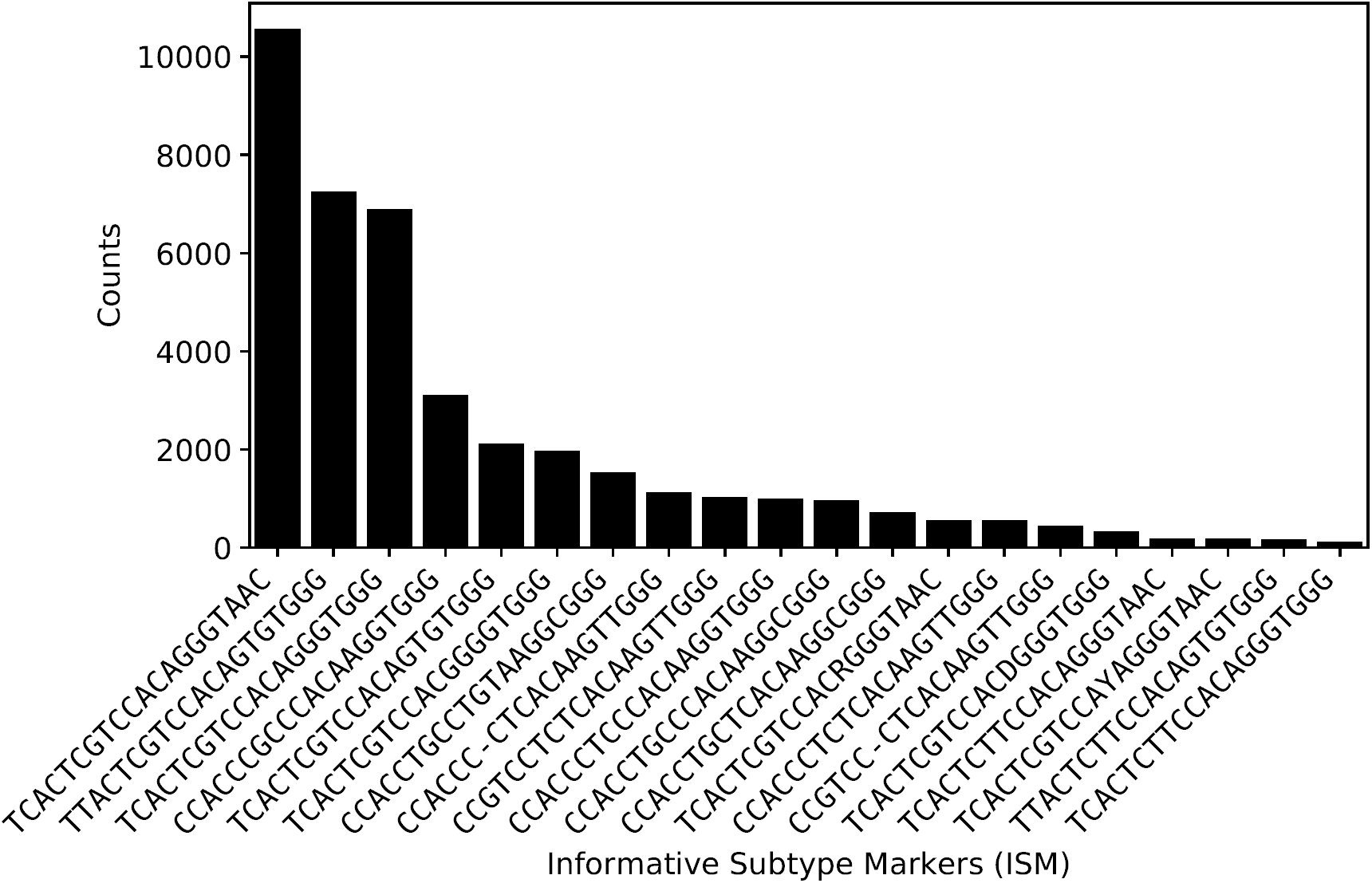
Number of sequences containing the 20 most abundant ISMs (after error correction) within the total data set (out of 45535 sequences), indicating the rapid drop off in frequency after the first few most prevalent ISMs.

Some potential reasons for the rapid drop off in the frequency relative to the diversity of ISMs may include the following: *First*, since the virus is transmitting and expanding so quickly, and the pandemic is still at a relatively early stage, there has not been enough time for mutations that would affect the ISM to occur and take root. In that case, we would expect the number of significant ISMs to rise over time. *Second*, the population of publicly available sequences is biased to projects in which multiple patients in a cluster are sequenced at once: e.g., a group of travelers, a family group, or a group linked to a single spreading event (there are sequences from cruise vessels in the database). We expect that the impact of any such clustering will be diminished in time as more comprehensive sequencing efforts take place. *Third*, ISMs may be constrained by the fact that certain mutations may result in a phenotypic change that may be selected against. In this case, we may expect a steep change in a particular ISM or close relative in the event that there is selection pressure in favor of the corresponding variant phenotype. However, as of yet there has been no solid evidence of mutations within SARS-CoV-2 that are associated with selection pressure, i.e., as being more transmissible or evading antibodies, though studies do suggest the possibility [19, 47, 48].

Figure 2 also shows that despite the application of the error correction method detailed in the Methods section, some symbols representing ambiguously identified nucleotides, such as `S` and `D` still remain in the ISMs. These represent instances in which there was insufficient sequence information to fully resolve ambiguities. We expect that as the number of publicly available sequences increases, there will likely be additional samples that will allow resolution of base-call ambiguities. That said, it is possible that the ambiguity symbols in the ISMs reflect genomic regions or sites that are difficult to resolve using sequencing methods, in which case the ISMs will never fully resolve. Importantly, however, because of the application of the error correction algorithm, there are fewer spurious subtypes which are defined due to variants arising from sequencing errors, and all remaining ISMs are still usable as subtype identifiers.

#### Potential functional significance of ISM locations

After the informative nucleotide positions were identified, we then mapped those sites back to the annotated reference sequence for functional interpretation [13]. As a practical matter, because the ISM is made up of the high-diversity sites within the SARS-CoV-2 genome, it inherently includes the major loci of genetic changes that are being identified in population studies worldwide. The ISM also excludes sites at the ends of the genome in which variation is most likely to be the result of sequencing artifacts. As shown in Table 1, we found that all but one of the nucleotide positions that we identified were located in coding regions of the reference sequence. The majority of the remaining sites (12/19) were found in the *ORF1ab* polyprotein, which encodes a polyprotein replicase complex that is cleaved to form nonstructural proteins that are used as RNA polymerase (i.e., synthesis) machinery [49]. One site is located in the reading frame encoding the S surface glycoprotein, which is responsible for viral entry and antigenicity, and thus represents an important target for understanding the immune response, identifying antiviral therapeutics, and vaccine design [50, 51]. High-entropy nucleotide positions were also found in the nucleocapsid formation protein, which is important for packaging the viral RNA [52]. A study has also shown that, like the spike protein, the internal nucleoprotein of the virus is significant in modulating the antibody response [53]. Other sites were found in the ORF3a and ORF8, which, based on structural homology analysis do not have known functional domains or motifs, and have diverged substantially from other SARS-related variants which contained domains linked to increased inflammatory responses [54, 55].

**Table 1.** Mapping ISM sites to the reference viral genome.

| Site | Nucleotide Position | Entropy | Annotation |
| --- | --- | --- | --- |
| 1 | 241 | 0.86092 | Non-coding Region |
| 2 | 1059 | 0.68346 | ORF1ab |
| 3 | 2480 | 0.23123 | ORF1ab |
| 4 | 2558 | 0.24007 | ORF1ab |
| 5 | 3037 | 0.86980 | ORF1ab |
| 6 | 8782 | 0.40949 | ORF1ab |
| 7 | 11083 | 0.45757 | ORF1ab |
| 8 | 14408 | 0.86807 | ORF1ab |
| 9 | 14805 | 0.49255 | ORF1ab |
| 10 | 17747 | 0.23734 | ORF1ab |
| 11 | 17858 | 0.23409 | ORF1ab |
| 12 | 18060 | 0.24005 | ORF1ab |
| 13 | 20268 | 0.28748 | ORF1ab |
| 14 | 23403 | 0.86829 | S surface glycoprotein |
| 15 | 25563 | 0.78760 | ORF3a |
| 16 | 26144 | 0.44564 | ORF3a |
| 17 | 28144 | 0.40928 | ORF8 |
| 18 | 28881 | 0.88582 | nucleocapsid phosphoprotein |
| 19 | 28882 | 0.88370 | nucleocapsid phosphoprotein |
| 20 | 28883 | 0.88174 | nucleocapsid phosphoprotein |

In sum, the majority of high-entropy sites are in regions of the genome that may be significant for disease progression, as well as the design of vaccines and therapeutics. Accordingly, ISMs derived from the corresponding nucleotide positions can be used for viral subtyping for clinical applications, such as identifying variants with different therapeutic responses or patient outcomes, or for tracking variation that may reduce the effectiveness of potential vaccine candidates. Unlike phylogenetic clusters, the ISM includes information about the single nucleotide variation (SNV) directly in the nomenclature. The subtypes which are identified are not a function of a selected clustering algorithm or a feature that has been selected as being relevant to a cluster.

#### Evaluating artifacts in ISM abundance due to error correction

Even though the SARS-CoV-2 data set appears to be large, it represents only a small sample of the full scope of cases. Therefore, tracking the pandemic requires using as much global data as possible, which means that imperfect sequence data must be tolerated to avoid losing potentially relevant samples. However, error correction will only be useful if it can maintain the integrity of the data, in particular, permit accurate identification of viral subtype abundance. In our case, we expect and do observe that the abundance of non-ambiguous ISMs will be inflated sightly by the error correction process. Supplementary file 3 — ISM inflation by error correction shows the inflation rate of highest-abundance ISMs after error correction.

We can see from Supplementary file 3 — ISM inflation by error correction that the error correction process only inflates the frequency of the highest-abundance ISMs in our database by less than 10%. To demonstrate that the error correction is a conservative process, we further show an ISM, CCACCCGCCCACAAGGTGGG, which is inflated by 10.79% as a case study. Half of the inflation arises due to sequences with ISM CCACCCGCCCACNAGGTGGG with 1.0 hamming distance away from the corrected ISM (there is an N instead of an A at position 13 in the ISM). Our error correction process corrects that `N` to `A` because all non-ambiguous ISMs with the same nucleotide configuration except for position 13 (non-ambiguous ISMs have an `A` at position 13 instead of `N`). Accordingly, ISM abundance inflation due to error correction will be generally conservative, and will not confound population-level analyses of ISM subtypes based on their relative abundance.

#### Sensitivity of ISM labels to the selection of the entropy threshold

To demonstrate the influence of the entropy threshold on ISM identification, we show a Sankey diagram in Figure 3. Figure 3 was constructed by first defining different sets of ISMs based on entropy threshold of 0.1, 0.23 (the major entropy threshold in our manuscript that result in 20 ISM sites), 0.4, 0.6 and 0.8. A stripe on the diagram represents an ISM (as labeled on the plot). The diagram tracks ISM identification according to sequences grouped by respective ISMs. For example, ISMs defined at a higher entropy threshold each likely identify more sequences (i.e. more sequences will likely have the same given ISM). Correspondingly, there will be an increased number of ISMs which are more refined (each identifying smaller collections of sequences), at a lower entropy threshold. The width of the stripe corresponds to the number of sequences with that ISM.

**Figure 3.**
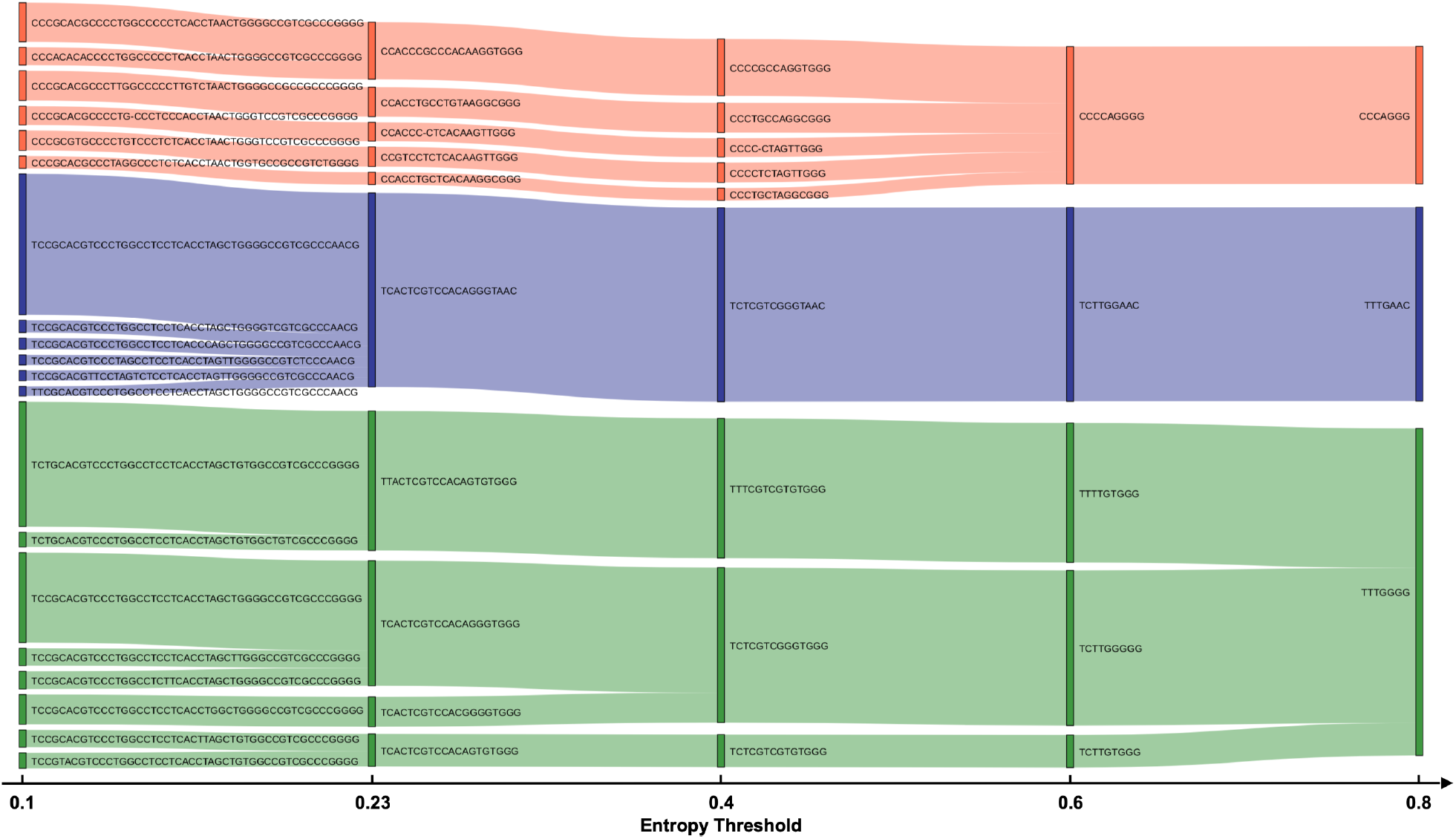
Sankey Diagram of the most abundant 20 ISMs defined by an entropy threshold of 0.1 and how they relate to ISMs defined at other entropy thresholds. This figure shows that the most abundant ISMs are generally stable for an entropy threshold between 0.23 and 0.4.

The Sankey diagram further shows that ISMs defined at a lower entropy threshold can “merge” together as the entropy threshold moves higher. For example, TCTTGGGGG and TCTTGTGGG are two different ISMs if we choose 0.6 as the entropy threshold. They can be differentiated by the 6th position (a `G/T` variation). However, when the threshold moves higher to 0.8, this position is dropped from the ISM, as its entropy now falls below the threshold. As a result, the two ISMs are merged into `TTTGGGG`. Some ISMs are stably identified throughout, while other ISMs merge together at different entropy thresholds. We can see from Figure 3 that the entropy threshold acts as a way to tune the resolution of subtype definition. When choosing a high entropy, positions that can differentiate relatively smaller (less abundant) subtypes are ignored. On the other hand, setting the entropy threshold lower reveals more ISM subtypes. For example, TCACTCGTCCACAGGGTAAC is defined at 0.23 threshold and when set to 0.1 threshold, 5 additional less-abundant ISMs emerge. We can thus observe, based on the diagram, that some subtypes are more “stable” markers than others. However, there are also some ISMs that are not sensitive to the selection of entropy threshold. For example, the subtype labeled as TTACTCGTCCACAGTGTGGG (particularly found in genetic sequences from New York state and some European countries, as discussed below) does not merge with other high-abundance ISMs until the entropy threshold is set to 0.7. Therefore, this ISM may be considered to be a stable marker. Overall, the most abundant ISMs are generally stable for an entropy threshold between 0.23 and 0.4.

We can also track how the entropy at individual variable positions evolve as a function of time. Figure 4 shows how the entropy at sites labeled by their position on the reference genome changes over time, as more sequences are collected and the genetic sequences change. Here, we visualize the dynamic of entropy values at different positions over time. That is, given a nucleotide position in the reference genome and the number of weeks since December 24, 2019, we compute the entropy of that position using all sequences collected by that time.

**Figure 4.**
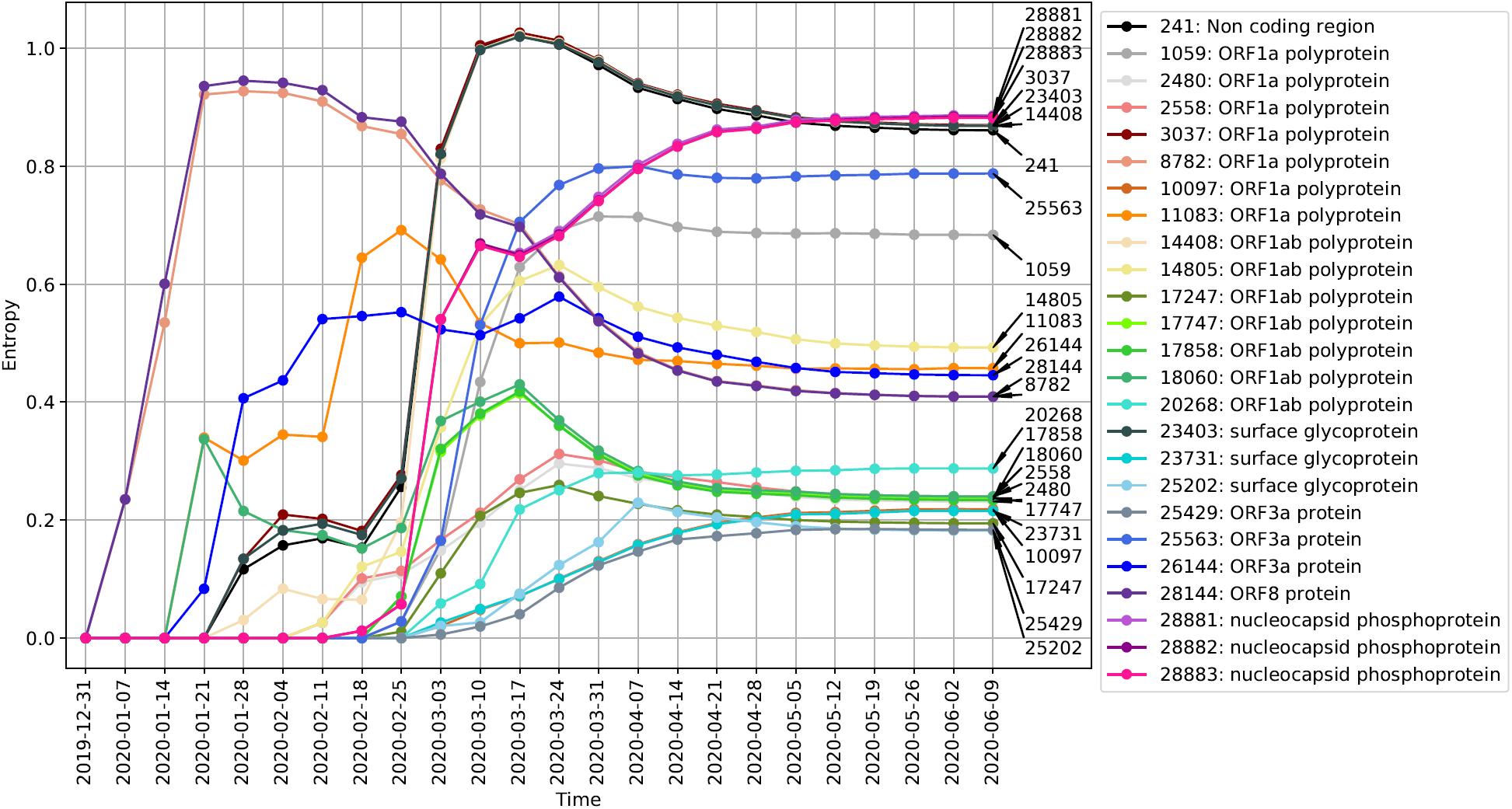
Entropy value changes over time at different ISM positions. Each curve in this figure represents the entropy of a highly variable position over time. The position in reference genome is labeled on the right end of the curve. The gene name associated with that position is labeled in the legend.

We can see that at the earliest stages of the pandemic, the ISM positions corresponding to nucleotide positions 8782 and 28144 in the reference sequence had the highest entropy, i.e., high variation in these two positions were found in sequences collected by Early February. Subsequently, the entropy values of these two positions drop. Notably, Figure 4 shows how the highly varying spike protein mutation at position 23403 (the `A` to `G` spike protein mutation, which has been found to be abundant in Europe and US and has since spread around the world [47], evidently became prevalent in the middle stage (in early March). In addition, we observe that there are some ISM positions which appear to covary, as indicated by the correlation between the changes in their entropy values. For example, the entropy of positions 8782 and 28144 covary, which is consistent with the correlation of the SNVs at these positions in genome sequence data available early on in the pandemic (i.e., before February 2020) [12].

#### Correlated/Covarying positions allow compression of the ISM representation

As indicated by Figure 4, the number of sites with sufficient entropy to be included in the ISM increases over time, as genetic changes occur and accumulate. This means that a representative ISM is a 20-base long identifier as of June 2020, which is unwieldy as a subtype identifier. Moreover, as shown in Figure 4, there are nucleotide sites with entropy covarying over time, representing correlations in genetic changes which result in redundancy in the ISM. Therefore, we may select a subset of positions to represent all the covarying positions to reduce the size of our ISMs. This results in a more compact identifier, which, at a small cost to resolution, provides for more efficient subtype differentiation and categorization.

Table 2 shows the most abundant nucleotide configurations at certain covarying positions and how many variations can be preserved after compression. The most abundant nucleotide configurations cover at least 96% of the sequences for each covarying group (the third column in Table 2). We further validate the groups of covarying nucleotide sites identified by the temporal entropy curve in Figure 4 by Linkage Disequilibrium (LD) analysis, which measures the degree of nonrandom association between two loci on a genome [56]. The results, included as Supplementary file 4 — Pairwise Linkage disequilibrium between high linkage sites, show that the sites within each covarying group have significant linkage with high pairwise *r*^2^ values (generally greater than 0.95). Linkage disequilibrium is a measure of the degree of nonrandom association between two loci [56]. This is in line with previous studies of LD on SARS-CoV-2 [12, 57, 58], which found, e.g., that positions 8782 and 28144 showed high significant linkage, with an *r*^2^ value of 0.954 [12].

**Table 2.** The most abundant nucleotide configurations at certain covarying positions and the number of sequences associated with them. The first column shows the group of covarying positions; The second column shows most abundant nucleotide configurations at those positions; The third column shows the number of sequences associated with the nucleotide configurations listed in the second column and the fourth column is the representative positions we picked to represent all the positions in this group. From this table, we can see that we can reduce the ISM size by 45% at a small cost to resolution

| Covarying group | NT configurations | Coverage | Representative position |
| --- | --- | --- | --- |
| 241, 3037, 14408, 23403 | TTTG, CCCA | 96.19% | 23403 |
| 2480, 2558 | AC, GT | 98.68% | 2558 |
| 8782, 28144 | CT, TC | 98.52% | 8782 |
| 17747, 17858, 18060 | CAC, TGT | 98.38% | 18060 |
| 28881, 28882, 28883 | GGG, AAC | 98.94% | 28881 |

We then select the representative positions with the highest entropy within each covarying group that can cover all of the most abundant nucleotide configurations. Compression reduces the ISM length from 20 to 11 nucleotides.

Table 3 shows the mapping between the original 20-NT ISM and 11-NT compressed ISM for ISMs associated with more than 100 sequences in the database. From the table we can see that most of the abundant 20-NT ISMs are assigned to unique 11-NT compressed ISMs. However, there are three 11-NT compressed ISMs that correspond to multiple 20-NT ISMs. For example, 20-NT ISM TCACTCGTCCACAGGGTAAC (10,565 sequences) and TCRCTCGTCCACAGGGTAAC (112 sequences) are merged to CCCGCCAGGGA after compression. These two subtypes are differentiated by position 3 in the long ISM (`A/R`). As such, defining subtypes based on compressed ISM will result in the inflation of a few principal subtypes by an amount of around 1%. Compressed ISM subtypes, therefore, conserve the distribution of major ISM subtypes. A more compact and easy-to-use subtype nomenclature may thus be utilized to quantitatively assess the relative subtype abundance at the population level.

**Table 3.**
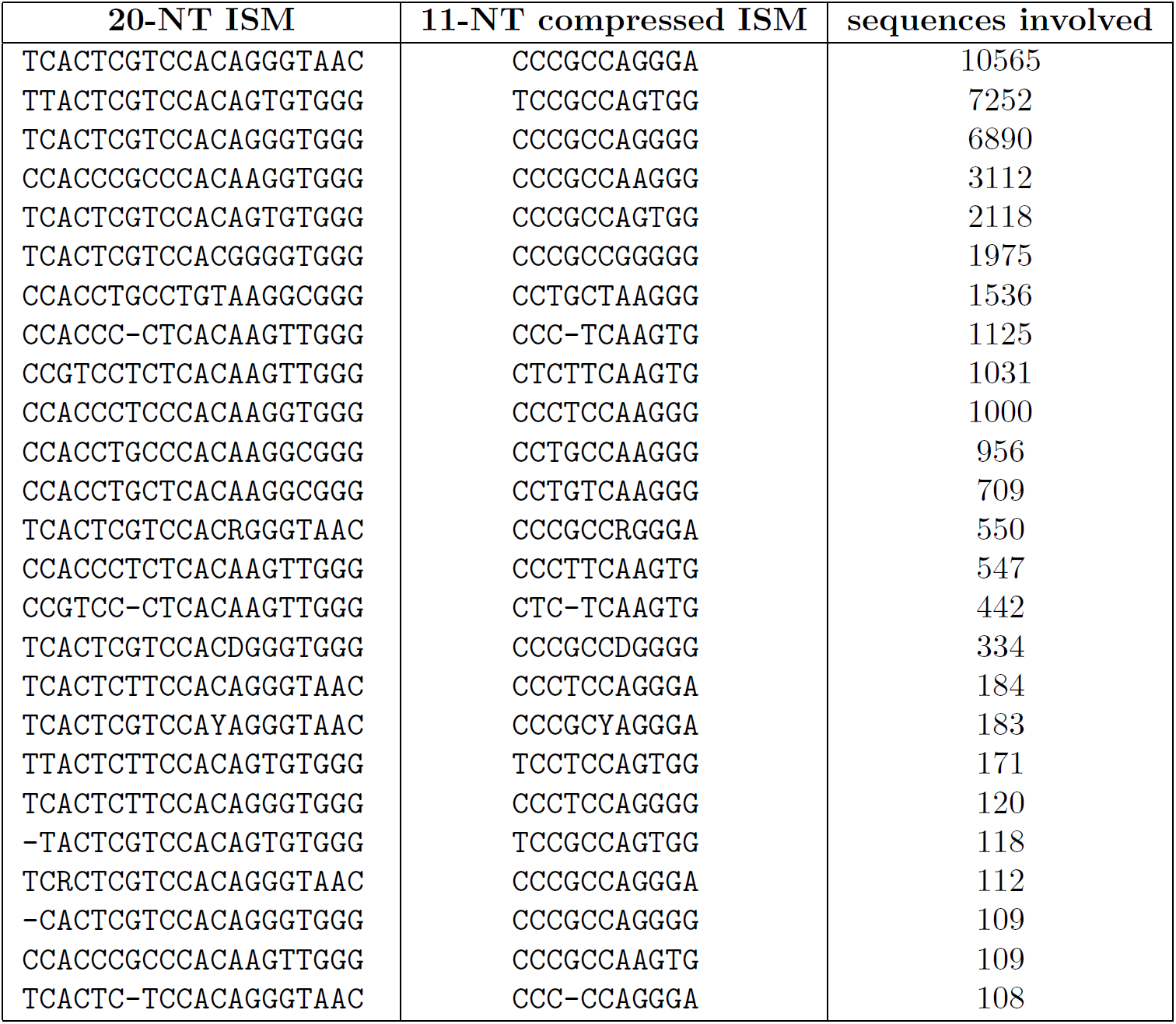
Map between 20-NT ISM and 11-NT compressed ISM.

### Geographic distribution of SARS-CoV-2 subtypes

To demonstrate that ISM subtypes can be used to analyze and visualize the spread of the SARS-CoV-2 pandemic, we describe the geographic distribution of the relative abundance of subtypes in different countries/regions worldwide, as well as in different states within the United States. Not only does this provide an illustration of the method’s capabilities, but it also permits comparison of the subtyping analysis with theories regarding viral spread between regions. Figure 5 shows the distribution of ISMs, each indicating a different subtype, in the regions with the relatively larger amount of available sequenced genomes. As shown therein, the ISMs are able to successfully identify and label viral subtypes that produce distinct patterns of distribution in different countries/regions. Beginning with Mainland China, the source of SARS-CoV-2 reference genome [13, 59] (NCBI Accession number: NC 045512.2), we observe two dominant subtypes, as indicated by relative abundance of the ISM among available sequences: CCCGCCAAGGG (as indicated on the plot, first seen on December 24, 2019, in sequences from Mainland China in the dataset), CCTGCCAAGGG (first seen on January 5, 2020 in sequences from Mainland China in the dataset).^7^

**Figure 5.**
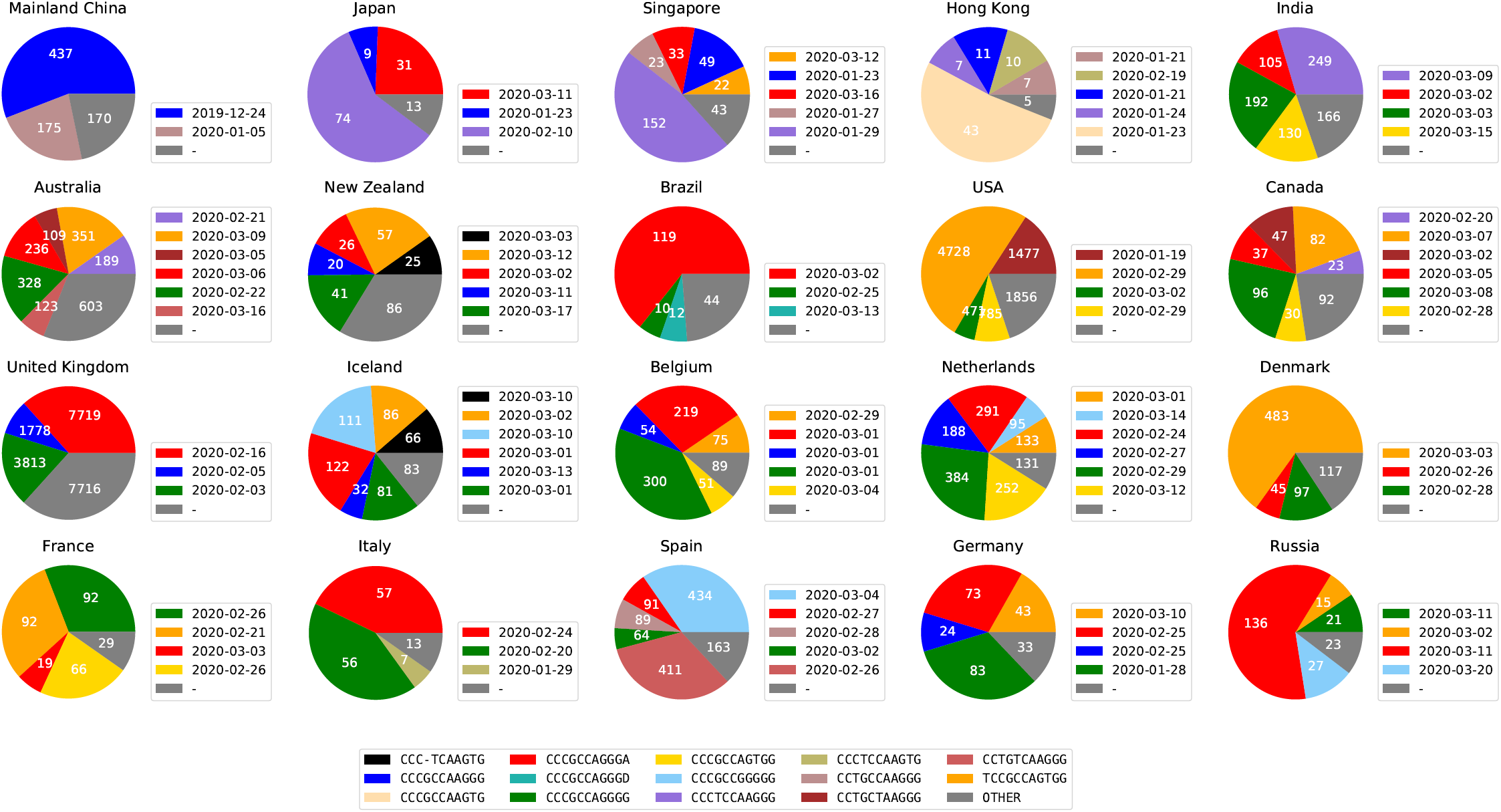
Major subtypes in countries/regions with the most sequences (in the legend next to each country/region, we show the date when a major subtype was first sequenced in that country/region). Subtypes with less than 5% abundance are plotted as “OTHER”. The raw counts for all ISMs in each country/region, as well as the date each ISM was first found in a sequence in that country/region, are provided in Supplementary file 6 — ISM abundance table of 20 countries/regions.

Another subtype, CCCTCCAAGGG, is the most abundant, i.e., dominant subtype in other Asian countries like Japan and Singapore (first detected on January 18, 2020, in sequences from Mainland China in the dataset). These subtypes are found in other countries/regions, but in distinct patterns, which may likely correspond to different patterns of transmission of the virus. Subtype CCCGCCAGGGA (first detected in February 16, 2020, in sequences from United Kingdom in the dataset) is found abundant in many European countries and then detected in Japan and Singapore later. This subtype has also been found in Canada and Brazil, suggesting a geographical commonality between cases in these diverse countries with the progression of the virus in Europe. Another prevalent subtype is TCCGCCAGTGG which is first detected in France in February 21, 2020. This subtype then becomes the dominant subtype in Denmark, USA, and one of the major subtypes in Canada and Germany. Both subtypes, CCCGCCAGGGA and TCCGCCAGTGG, have an A23403G mutation (corresponding to position 14 in the ISM) which has been discussed in recent studies [19, 47].

The data further indicate that the United States has a distinct pattern of dominant subtypes. In particular, the subtype with the highest relative abundance among U.S. sequences is CCTGCTAAGGG, first seen on February 20, 2020. This subtype has also emerged as one of the major subtypes in Canada, with the first sequence being found on March 5, 2020.

We also found that some states within the United States have substantially different subtype distributions. Figure 6 shows the predominant subtype distributions in the states with the most available sequences. The colors shown on the charts are also keyed to the colors used in Figure 5, which allows for the visualization of commonalities between the subregional subtypes in the United States and the subtypes distributed in other regions. It is obvious that east coast states and west coast states demonstrate different ISM distributions.

**Figure 6.**
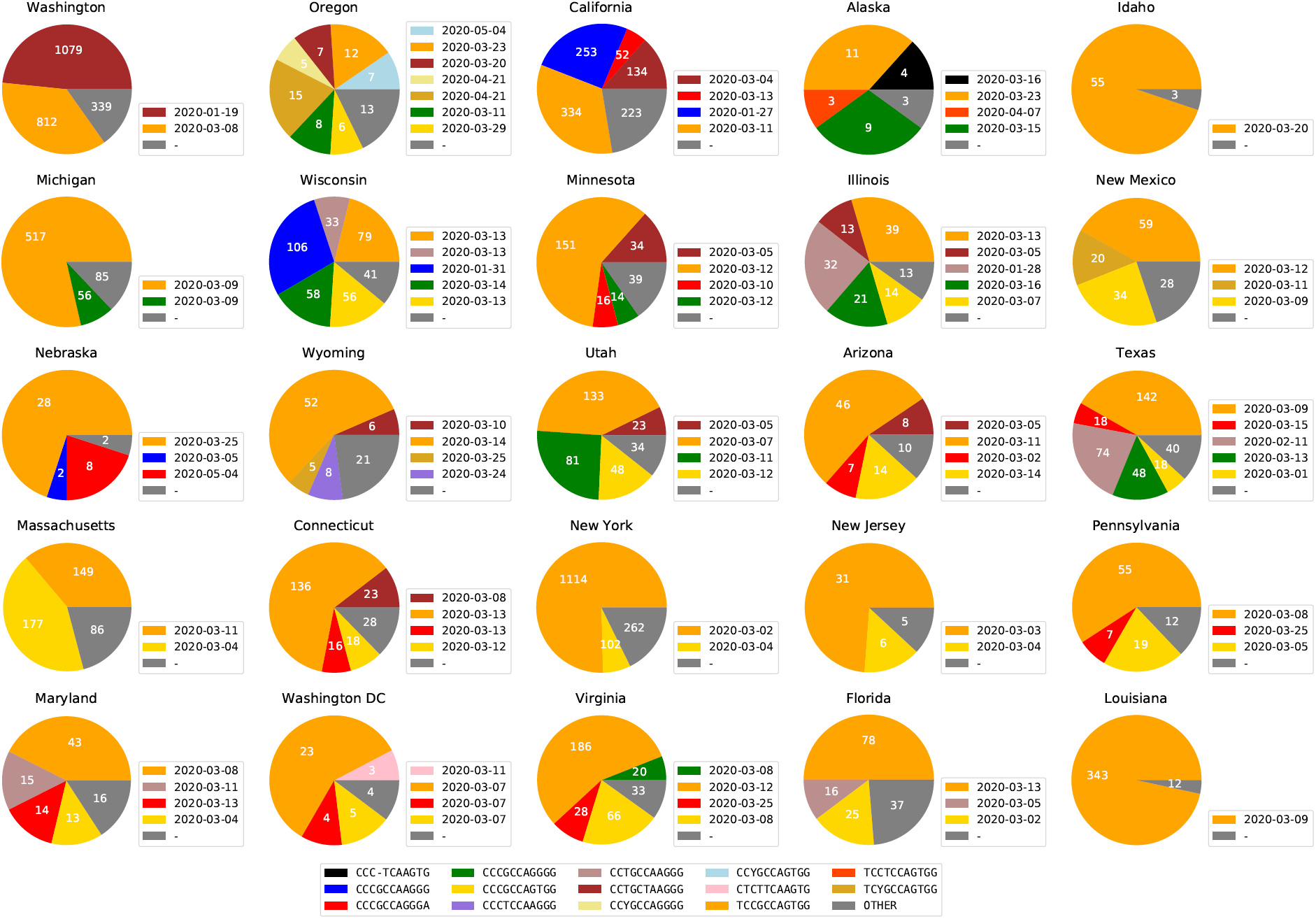
Viral subtype distribution in the United States, showing the 25 states with the most sequence submissions. Subtypes with less than 5% abundance are plotted as “OTHER”. The raw counts for all ISMs in each state, as well as the date each ISM was first found in a sequence in that state, are provided in Supplementary file 7 — ISM abundance table of 25 U.S. states

Most prominently, New York is dominated by a subtype, TCCGCCAGTGG, which is also highly abundant among sequences from European countries, including France, Denmark, Germany, and Iceland. California, on the other hand, includes as a major subtype, CCCGCCAAGGG, which is also a major subtype in Mainland China, as shown in Figure 5. The most abundant subtype in Washington, CCTGCTAAGGG, is also a major subtype in other states in United States. This CCTGCTAAGGG subtype is also found in substantial abundance in Canada as well. This is consistent with the hypothesis that this subtype is endogenous to the US.

Regions with similar genetic variant patterns are identifiable in Figure 5, but only at a qualitative level. As described in the Methods section, the ISM abundance table can be used to provide a quantitative analysis of the similarity between the genetic variation patterns of countries and regions. Figure 7 shows a visualization of the difference in genetic subtype patterns between different countries and regions using Principle Components Analysis (PCA) as described in the Methods, projecting onto the first two principle components. We can see that a few European countries form small clusters of similar ISM abundance, i.e., a similar subtype distribution. This implies that similar SARS-CoV-2 subtypes are shared by these countries, e.g., Austria, Netherlands, Germany, and Sweden. Most Asian countries are projected to the upper right part of the PCA plot, in contrast to North American countries and a few European countries clustered towards the bottom. This indicates the difference between the dominant genetic subtypes of ISM and patterns of genetic variation between these regions of the world. In particular, the separation in this ISM subtype space futher supports the hypothesis that the outbreak in New York is linked to some travel cases from European countries, such as France. To further validate the utility of ISMs for subtyping, we show an analysis of the geographical distribution of the dominant subtypes in Italy in Supplementary file 8 — Geographical distribution of the dominant subtypes in Italy.

**Figure 7.**
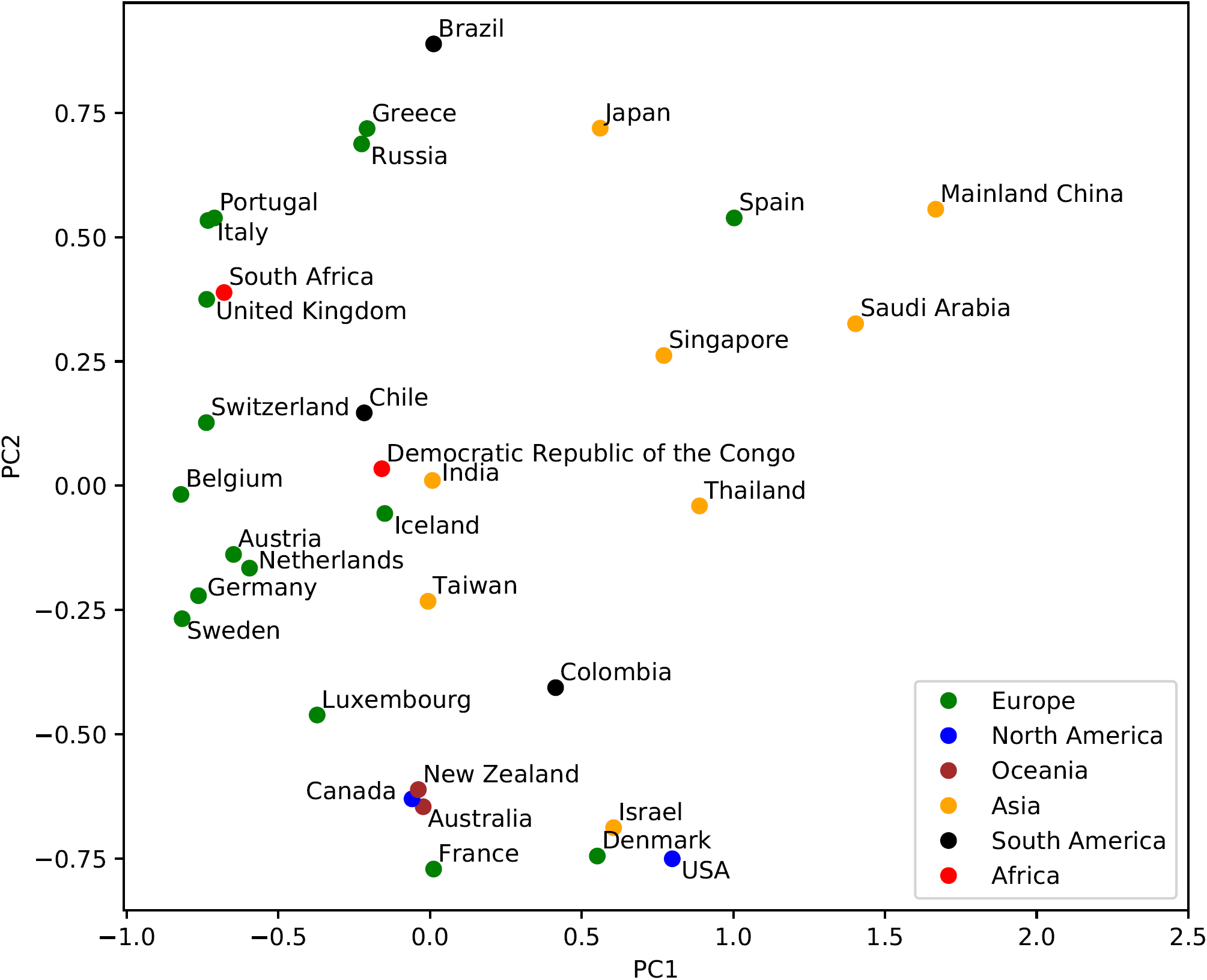
Country/region-specific patterns of viral genetic variation visualized by the first two principle components of the Bray-Curtis dissimilarity matrix. The regions are color coded by continents. Each point represents the SARS-CoV-2 genetic variation pattern of the labeled country/region based on the abundance of different ISMs in the country/region.

### Temporal dynamics of SARS-CoV-2 subtypes

The present-time geographical distributions shown in Figure 5, 6, and 7 suggest that ISM subtyping may identify the temporal trends underlying the expansion of SARS-CoV-2 virus and the COVID-19 pandemic. To demonstrate the feasibility of modeling the temporal dynamics of the virus, we first analyzed the temporal progression of different ISMs on a country-by-country basis. This allows examination of the complex behavior of subtypes as infections expand in each country and the potential influence on regional outbreaks by subtypes imported from other regions.

As described in the Methods section, we graph how viral subtypes are emerging and growing over time, by plotting the relative abundance of viral subtypes in a country/region (via the most frequently occurring ISMs over time) in Figure 8 and Figure 9. As discussed above, through the pipeline we have developed, these plots use a consistent set of colors to indicate different ISMs (and are also consistent with the coloring scheme in Figure 5).

**Figure 8.**
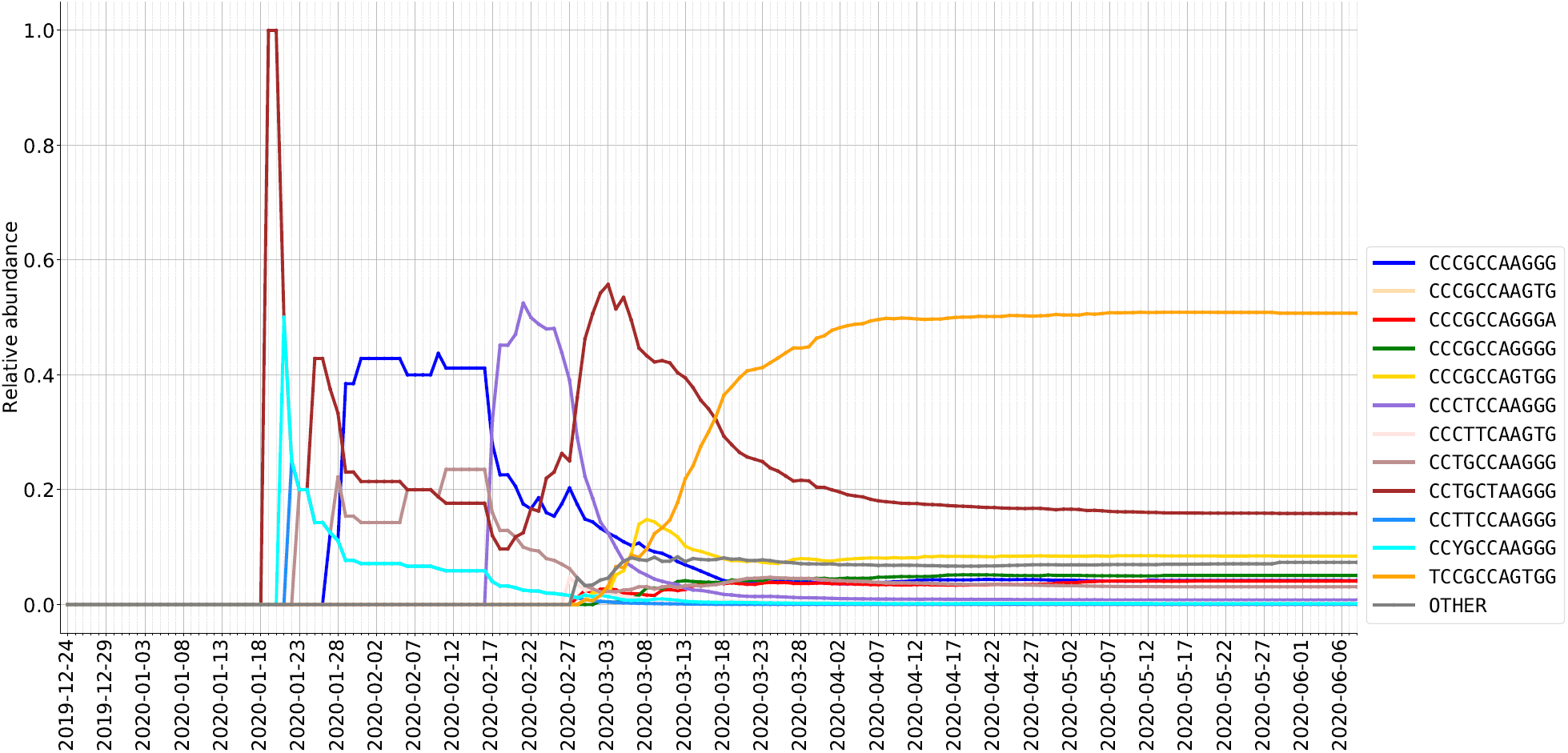
Relative abundance (%) of ISMs in DNA sequences from USA as sampled over time.

**Figure 9.**
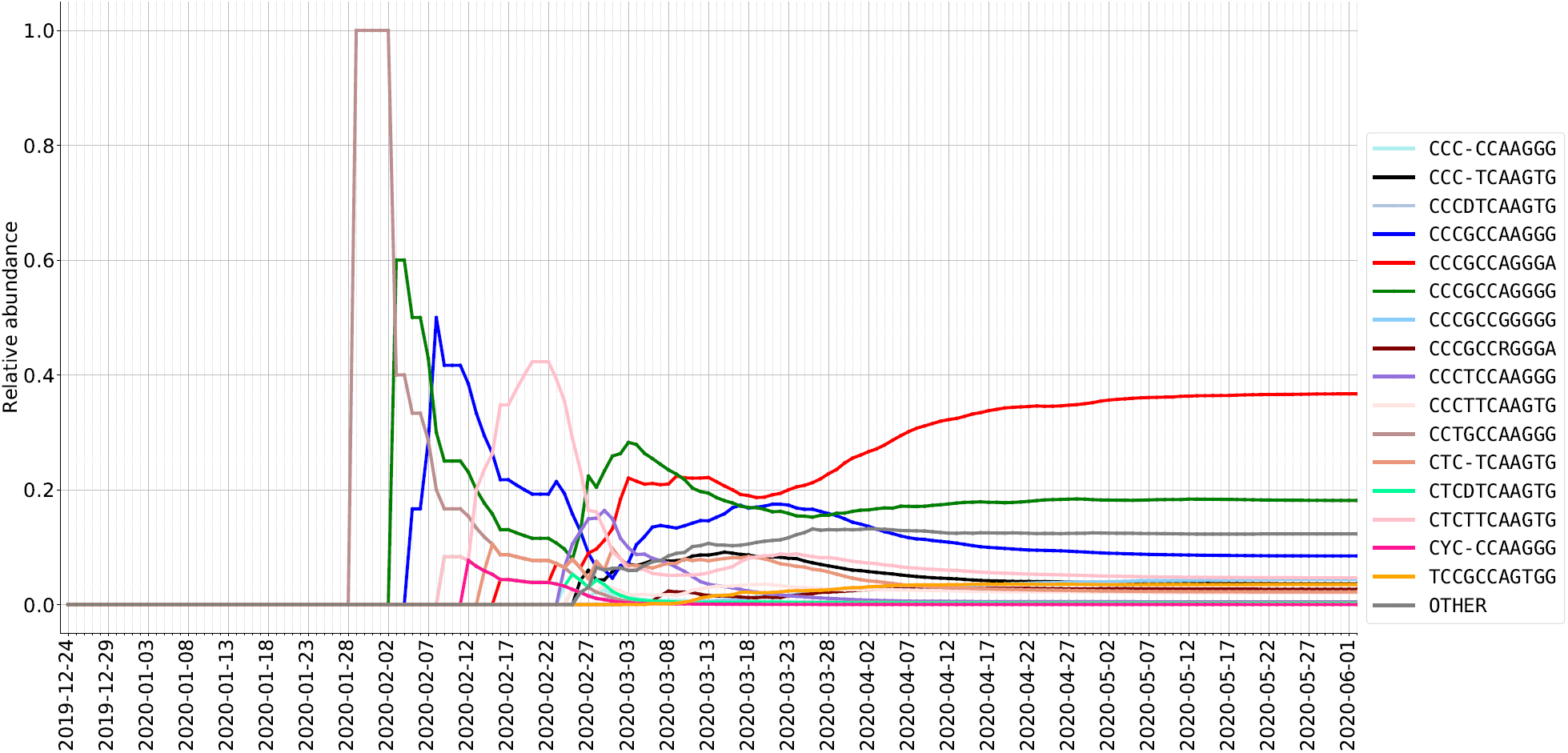
Relative abundance (%) of ISMs in DNA sequences from the United Kingdom as sampled over time.

In the United States, we can observe a few waves of different subtypes. For example, in the early stage (late January and early February), the predominant subtype is the same as that of Mainland China. In contrast, the most abundantly found subtype in late February and March, CCTGCTAAGGG, is not abundant in either Asia or Europe. However, this subtype has been found in a substantial number of sequences in both Canada and Australia. It is plausible, therefore, that the CCTGCTAAGGG subtype has become abundant as the result of community transmission in the United States, and has been exported from the United States to these other countries. Interestingly, while the CCTGCTAAGGG subtype has been found across the United States, as shown in Figure 6, it has not been found to be substantially abundant in New York. Over time, however, within the United States the dominant subtype has become TCCGCCAGTGG, which is the predominant subtype in New York state (and linked to the dominant subtype in many European countries).

As shown in Figure 9 and additional temporal plots for the Netherlands and Spain contained in the Supplementary Material, the subtype distribution in sequences within European countries differs significantly from that of North America and Australia. In particular, as detailed above, the European dynamics of SARS-CoV-2 appear to reflect the theory that in many European countries, initial cases may have been due to travel exposure from Italy, rather than directly from China. For instance, we observe that the United Kingdom data shows the same early subtypes as those of Mainland China which were also observed in Australia and Canada, i.e., CCTGCCAAGGG and CCCGCCAAGGG. The CCCGCCAAGGG subtype emerged as a highly abundant subtype in United Kingdom data in early February. This subtype was also been found with great frequency in the Netherlands and Australia, but not in Spain, suggesting additional viral genetic diversity within Europe for further study.

All inferences drawn from observed temporal trends in subtypes based on the genome sequence dataset—whether based on ISM or phylogeny-based methods–will be limited by important caveats, including: 1) The collection date of the viral sequence is usually later than the date that the individual was actually infected by the virus. Many of those individuals will be tested after they develop symptoms, which may only begin to arise several days or even two weeks after infection according to current estimates [60]. 2) The depth of sequencing within different regions is highly variable. As an extreme case, Iceland, which has a small population, has 1.3% of all sequences in the complete data set. Italy, on the other hand, had a large and early outbreak but has disproportionately less sequencing coverage (133 sequences).

### Evaluating the ability of ISM-defined subtypes to track significant genetic changes during the SARS-CoV-2 pandemic

In our results section, we identified a few widespread ISM subtypes, e.g., TCCGCCAGTGG that dominates New York and some ISM subtypes that are unique to a region, e.g., CCTGCTAAGGG that is mostly found in North America. In this section, we show related literature and how their results relate to ours. We primarily use the original 20-nt ISM identifiers in this section, rather than the compressed ISM, in order to discuss all the positions identified by our entropy analysis and relate them to the literature.

Subtype prevalent in New York and some European countries TTACTCGTCCACAGTGTGGG (TCCGCCAGTGG in compessed form)

This subtype has been dominating the US since mid-March, as shown in Figure 8. In Figure 6, we can see that this subtype dominates many states including New York (first seen early March in New York). Additionally, as shown in Figure 5, this same subtype has been dominant in European countries, first observed in sequencing data in late February. The first detection dates in New York (later) and Europe (earlier) align with the hypothesis of European travel exposure being the major contributor to the New York outbreak of SARS-CoV-2. Various studies have demonstrated the SNV C14408T in ORF1b to be associated with a virus subtype found abundantly in New York as well as multiple European countries [16, 61, 62], which is designated as an ISM hotspot site 8 in Table 1. These studies also identified a SNV of A23403G in the S spike protein to be heavily associated with the dominant subtype of both Europe and New York, correlating to ISM hotspot site 14 from our analysis. Our temporal entropy plot in Figure 4 further indicates that these two sites are covarying. Lastly, the studies also reported a SNV of G26144T, which corresponds to ISM site 16 and has been observed in the predominant subtypes found in Europe and New York.

#### Subtype potentially endogenous to the United States CCACCTGCCTGTAAGGCGGG (CCTGCTAAGGG in short form)

This is the prevalent subtype characteristic within Washington state through the lastest update of the sequencing database analyzed in this paper (June 2020). It has been linked to the endogenous spread of the virus across the United States [18, 63]. According to our ISM analysis, this subtype is separated by a hamming distance of 3 from one of the major subtype of the outbreak in China, CCACCTGCCCACAAGGCGGG (the differences are at ISM positions 10, 11 and 12). Viral spread is suspected to be due to primary exposure of an individual from China to Washington state, designating this case as “WA1” [17, 61]. “WA1” lineage is noted to have three characteristic SNVs, namely, C17747T, A17858G, and C18060T which correspond matches with our ISM positions of 10, 11 and 12 respectively [16–18, 61, 62]. While “WA1” is suspected as the primary subtype for viral spread in Washington state, there are cases that have shown additional SNVs, which suggest mutational variation from the “WA1” strain. These SNVs include C8782T and T28144C [16, 17, 61] and correspond to hotspot sites 6 and 17 respectively. The same major subtypes seen in Washington state were also identified in positive cases in Connecticut (also detected by Figure 6 using our ISM). It is highly probable that there was trans-coastal exposure due to domestic travel from Washington state into Connecticut, due to the various high-volume airports that are present in and around this state [18].

#### Subtypes including the A23403G/D614G spike protein variant

The SNV A23403G (resulting in D614G variant in spike protein) is a major viral mutation that has been observed in the major European countries of Italy, Spain, France, as well as Middle Eastern regions of Turkey and Israel [16, 64–66]. Some studies suggest that this D614G variant of the S spike protein provides greater survival and transmission ability to the virus, however there need to be additional studies conducted to confirm these claims [64]. This position corresponds to ISM position 14. Based on our ISM table, we can quickly navigate to this position and plot the abundance of different variants at this position over time. Figure 10 shows how the abundances of the variants at position 23403 change over time. We can quickly make this plot by indexing all the ISMs at position 14 and grouping them temporally. Indeed, Figure 10 illustrates how, in late February, A23403G started to take off in abundance and has quickly overwhelmed the initially more prevalent subtype.

**Figure 10.**
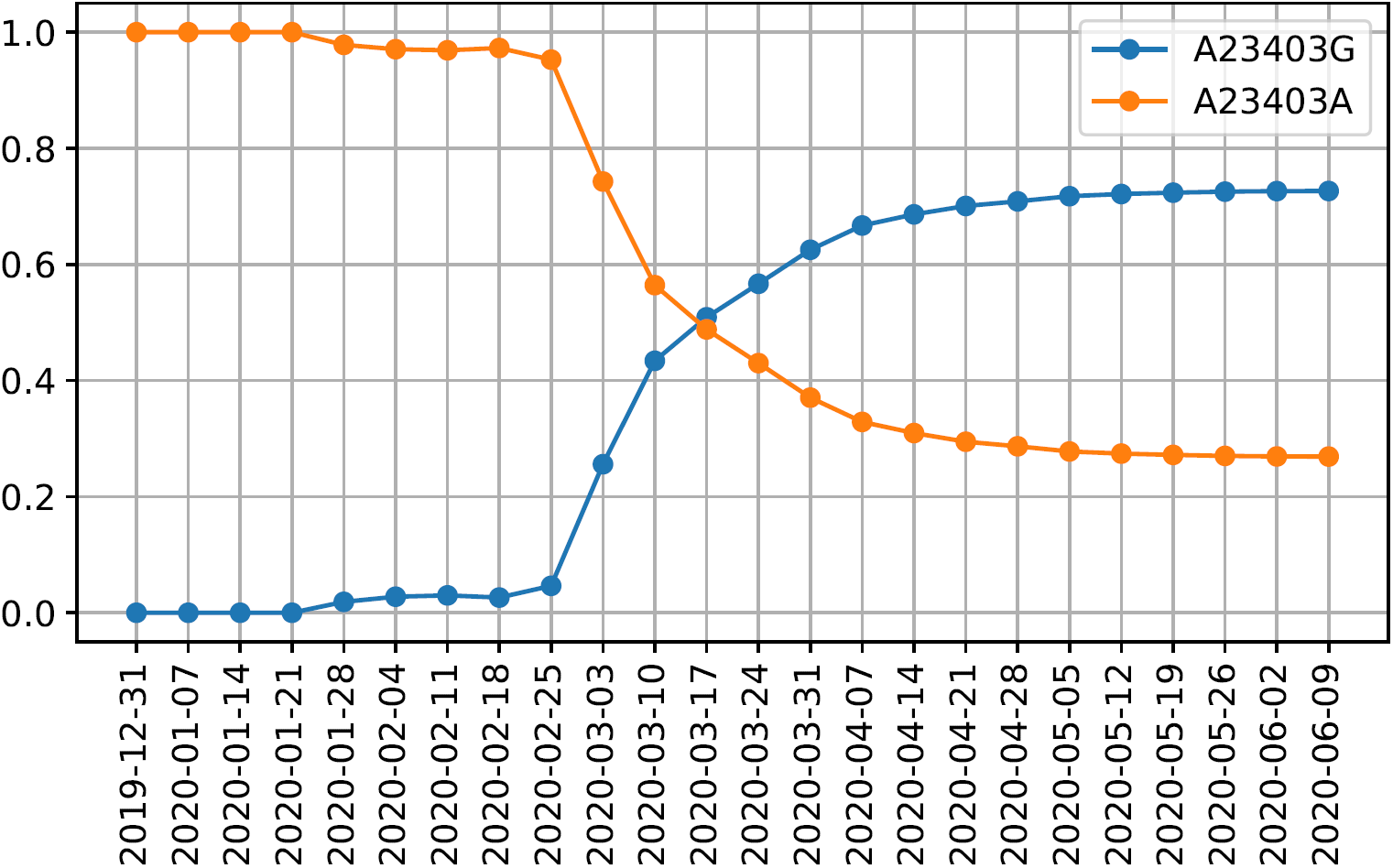
The relative abundance of variants of the D614G spike protein mutation (position 14 in our ISM and position 23403 in the reference genome).

### Comparison of ISM-defined subtypes to clades identified using phylogenetic trees

As discussed in the foregoing, subtypes defined by ISM are differentiated based on single nucleotide variants, which may eventually be found to represent functionally significant mutations in the viral genome. The ISM, however, does not include phylogenetic information, which sharply limits the utility of the ISM to infer patterns of viral evolution. Nevertheless, ISM-defined subtypes do correspond well with clusters of sequences based on phylogenic reconstruction. To identify whether the ISM may still be an effective identifier of genetic subtypes within the context of viral evolution, we compare subtype identification using the ISM and the phylogenetic tree structure. In particular by comparing the ISM-defined groups of sequences identified by our pipeline with the phylogenetic tree-based clusters (i.e., clades) identified by the Nextstrain group [1]. We do so by placing an ISM of interest at the lowest common ancestor (LCA) node of sequences containing that ISM on the phylogenetic tree produced by Nextstrain. Then, we compare the branch length between the root and LCA, which is considered as the evolutionary distance between a node and the reference sequence, and the Hamming distance between a given ISM and the reference ISM, CCACCCGCCCACAAGGTGGG (CCCGCCAAGGG in short form), which represents the degree of difference (by number of SNVs) between ISMs.

Figure 11 shows that Hamming distance (dark-colored) has a high correlation with the LCA branch length (gray-colored). This means that the Hamming distances between ISMs are able to consistently reflect evolutionary distance at a high level. There are a few outliers though; for example, CCACCTGCTCACAAGGCGGG (CCTGTCAAGGG in short form) has higher LCA branch length but lower ISM Hamming distance. This indicates that some evolutionary signals will be missed by grouping sequences by ISM, likely because the signals are contained in lower-entropy genomic regions which are unrepresented in the ISM. Conversely, we observe that the phylogenetic clades identified by Nextstrain are imperfect with respect to their preservation of SNV information. Nextstrain identifies the clades based on whether they contain at least two prevalent SNVs. But, presumably because the clades are identified by whole genome sequence clustering, not every sequence within a clade will necessarily include those SNVs.

**Figure 11.**
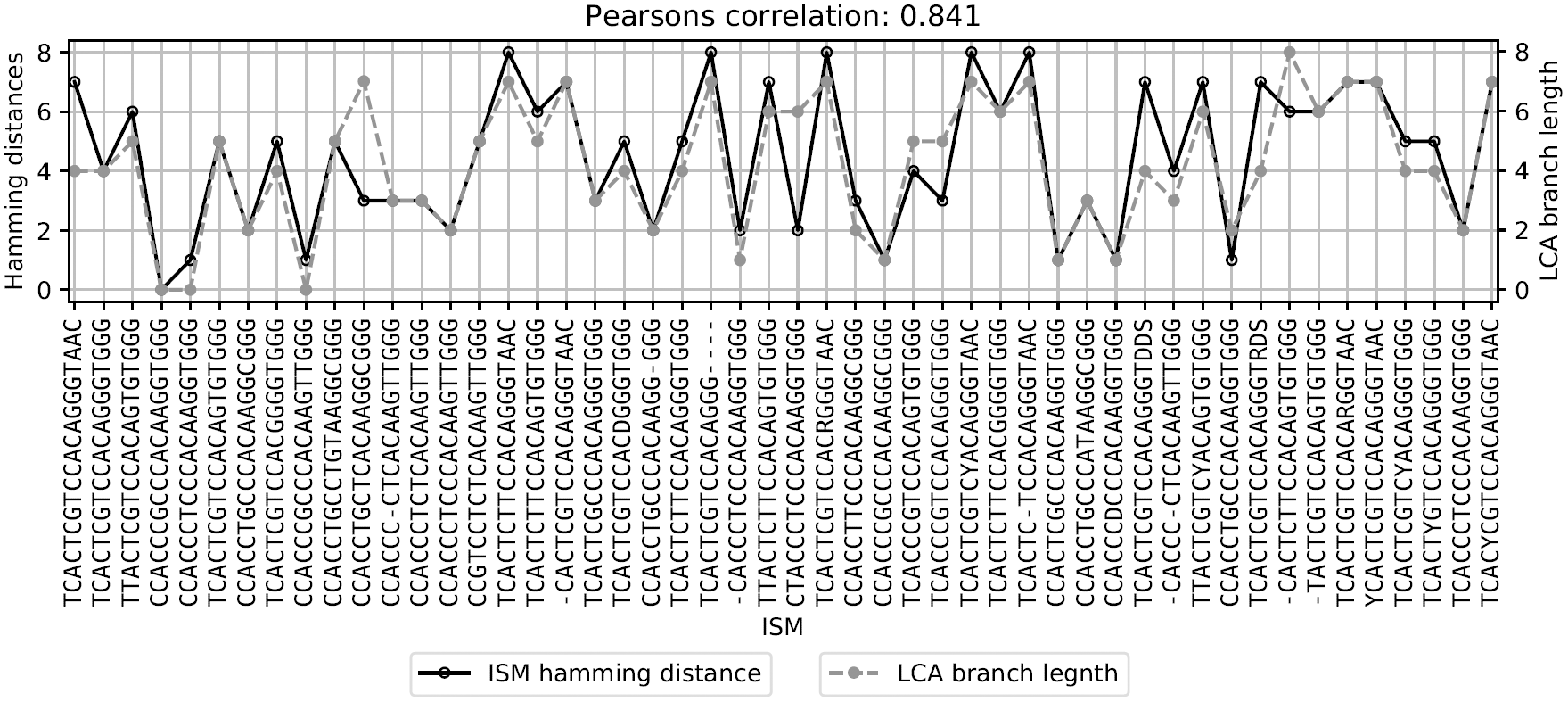
Demonstrating ISM distance is representative of the phylogenetic distance. Shown is the correlation between the branch length from LCA of sequences with an ISM of interest to the root and the Hamming distance between the ISM and the reference ISM, CCACCTGCTCACAAGGCGGG

Moreover, not only can the ISM pipeline effectively define meaningful viral subtypes, but it can also do so with greater computational efficiency than tree reconstruction methods. Fasttree [67], on its fastest setting, is reportedly the fastest tree reconstruction method (orders of magnitude faster than most machine learning methods). Fasttree theoretically executes at *O*(*N* ^1.25^ × *log*(*N*) × *L* × *a*) time, where *N* is the number of unique sequences, *L* is the width of the alignment, and *a* is the size of the alphabet. For Shannon’s entropy, the basis of ISM definition used in our work, the computation is *O*(*L* × *N* × *a*) where *L* is the number of loci and *N* is the number of sequences, and *a* is the size of the alphabet. Accordingly, the computational time required to enumerate subtypes using the ISM is substantially reduced, i.e., a function of the thresholded loci reduced and number of sequences instead. One caveat is that the ISM method requires multiple sequence alignment to identify high entropy sites, which can be a computationally intensive process. However, phylogenetic tree methods based on whole genome sequences require that as well. And, ISM identification may be done on new sequences using previous positions between multiple sequence alignment updates.

In sum, ISM can provide a compact and effective representation of a sequence as it includes the essential genetic variation information, while also including a substantial amount of molecular evolution information.

We further assess how ISMs defined at different entropy thresholds relate to the clades identified by Nextstrain. We compute homogeneity and completeness scores between ISM labels and Nextstrain clades as a function of entropy threshold. Homogeneity measures the extent to which ISMs each identify only sequences in one clade. Completeness measures the extent to which sequences that are members of a given clade are identified by a common ISM. Figure 12 shows the homogeneity and completeness as a function of the entropy threshold used to define ISMs. As shown therein, sequences with a common ISM are generally assigned to a common clade, and sequences from a given clades also often identified by a set of few ISMs. As the entropy threshold increases, ISMs correspondingly moving “upwards” through the phylogenetic tree to better represent a clade, increasing completeness while maintaining high homogeneity. Conversely, as the entropy threshold lowers, ISMs increase in their resolution, corresponding to an increase to almost perfect homogeneity but with low completeness.

**Figure 12.**
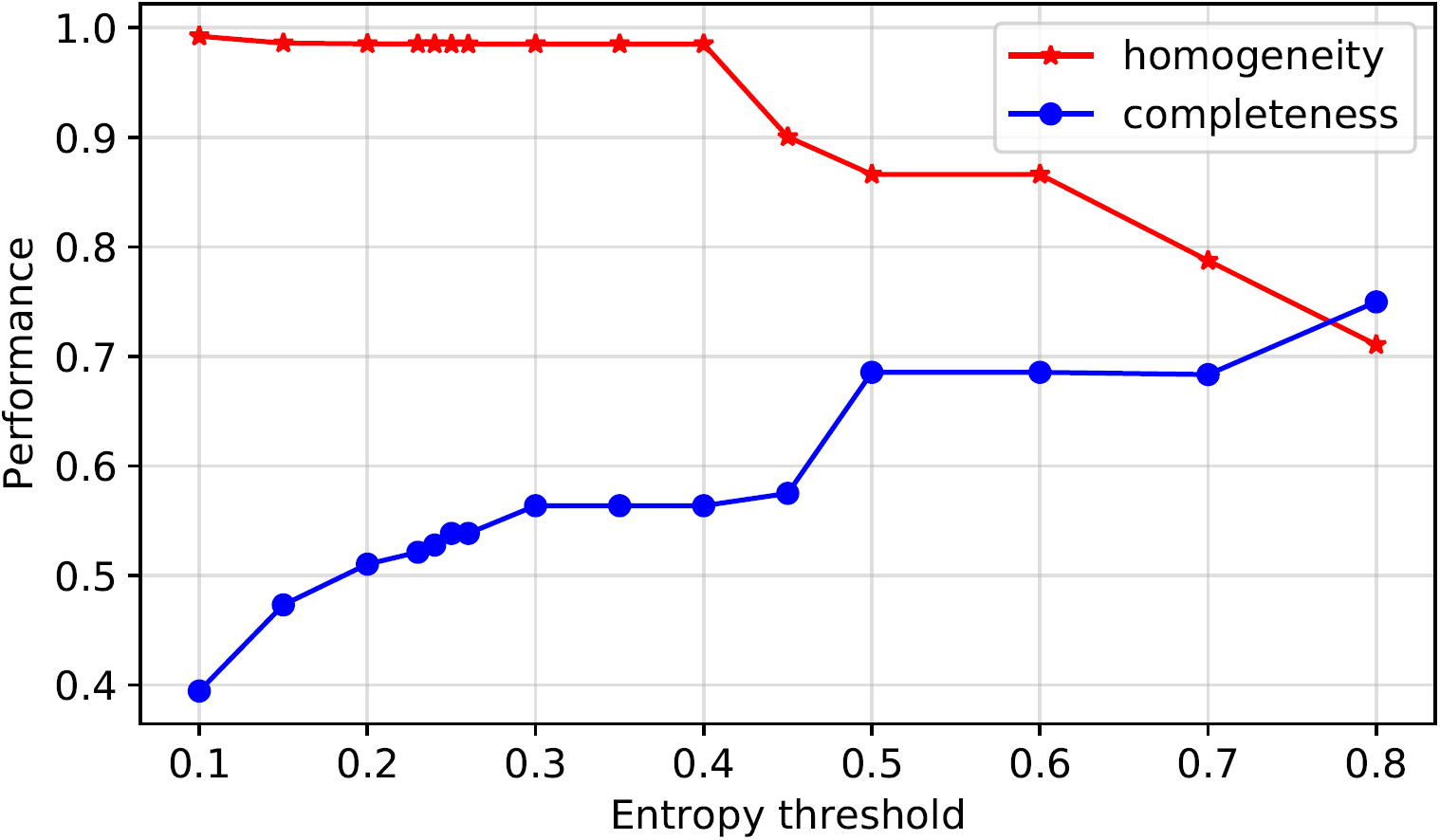
The effect of the entropy threshold on ISM membership to Nextstrain clade representation. At a low entropy threshold, each ISM contains sequences that nearly all belong to the same clade (high homogeneity) but the clade contains multiple ISMs (low completeness). As the entropy threshold rises, ISMs gain more sequences (some of which belong to other clades) and the clades contain fewer distinct ISMs. Thus, there is a trade-off with the entropy threshold, but the sweet spot is around 70-80% on both metrics, showing that ISMs capture some aspect of phylogenetics but have their own characteristics.

## Conclusions

In this paper, we present a pipeline for subtyping SARS-CoV-2 viral genomes based on short sets of highly informative nucleotide sites (ISMs). Our results demonstrate the following key features of ISM-based subtyping. *First*, the ISM of a sequence preserves important nucleotide positions that can help to resolve different SARS-CoV-2 subtypes. ISMs provide a quick and easy way to track key sets of SNVs which are *covarying* as the SARS-CoV-2 pandemic spreads throughout the world. The SNVs which consistently covary with the spike protein variant has rapidly become prevalent throughout the world and may be a potential link to increased viral transmission [4, 19, 47]. *Second*, ISM-based subtypes are able to capture the majority of phylogenetic relationships between viral genomes that are represented in *Nextstrain* tree clades. ISM analysis shows promise as a complement to phylogenetic classification, particularly given the limits of phylogenetics at early stages in the pandemic (e.g., due to uncertainty regarding key assumptions, such as the rate of the molecular clock and confidence in branches) – while also being more computationally efficient. *Third*, ISM subtyping can provide robust and informative insight regarding the geographic and temporal spread of the SARS-CoV-2 sequences, as well potentially be a way to identify phenotypic variants of the virus. For example, in this paper, we show that the distribution of ISMs is an indicator of the geographical distribution of the virus as predicted by the flow of the virus from China, the initial European outbreak in Italy, and subsequent development of local subtypes within individual European countries as well as interregional differences in viral outbreaks in the United States.

An important caveat of all viral analyses, including subtyping, is that they are limited by the number of viral sequences available. Small and/or non-uniform sampling of sequences within and across populations may not accurately reflect the true diversity and distribution of viral subtypes. However, the ISM-based approach has the advantage of being scalable as sequence information grows, and with more information, it will become both more accurate and precise for different geographic regions and within subpopulations.

Using ISM subtyping pipeline on continuously updated sequencing data, we are capable of updating subtypes as new sequences are identified and uploaded to global databases. We have made the pipeline and updated analyses available on Github at https://github.com/EESI/ISM and an interactive website at https://covid19-ism.coe.drexel.edu/. In the future, as more data becomes available, ISM-based subtyping can be employed on subpopulations within geographical regions, demographic groups, and groups of patients with different clinical outcomes. Efficient subtyping of the massive amount of SARS-CoV-2 sequence data will therefore enable quantitative modeling and machine learning methods to develop improved containment and potential therapeutic strategies against SARS-CoV-2. Moreover, the ISM-based subtyping scheme and associated downstream analyses for SARS-CoV-2 are directly applicable to other viruses, enabling efficient subtyping and real-time tracking of potential future viral pandemics.

## Acknowledgments

We downloaded all SARS-Cov-2 sequences available from and acknowledge the contributions of the Global Initiative on Sharing All Influenza Data (GISAID) EpiFlu database, which has made accessible novel coronavirus sequencing data, including from the NIH Genbank resource [6]. We would also like to acknowledge the authors, originating and submitting laboratories of the sequences from GISAID’s EpiFlu Database on which this research is based, as well as all future SARS-CoV-2 sequence contributors in GISAID’s EpiFlu Database. A list of contributors to the data used in this paper is included in Supplementary file 15 — Acknowledgements of sequences this research is based on. This work was partially supported by NSF grant #1919691. This work also used the Extreme Science and Engineering Discovery Environment (XSEDE), which is supported by National Science Foundation grant number #ACI-1548562, and Drexel’s University Research Computing Facility (URCF).

## Author Contributions

ZZ contributed to the conceptualization of the problem and solution, data curation, methodology, software, validation, visualization, and original draft preparation. BAS contributed to the conceptualization, data curation, methodology, software, validation, original draft preparation, and visualization. CM and KZ contributed to literature review and original draft preparation. GLR contributed to the project administration, conceptualization, methodology development, acquiring resources, validation, visualization, and original draft preparation.

## Supplementary Files

### Supplementary file 1 — Masked entropy threshold analysis

This figure shows the histogram of entropy and the frequency of N and - (Left: Histogram of masked entropy values of sites calculated without considering ambiguous bases or gaps (N’s and -’s). The red line demonstrates the high-entropy threshold used to define ISM sites (> 0.23); Right: Many genome positions in the alignments had nearly all ambiguous bases and/or gaps (the peak at around 1.0). On the other hand, the peak at around 0 represents high quality positions with a fewer number of N and - present. Sites with large number of N’s and - should be filtered out because a large number of N’s and -’s at a position is typically due to sequencing error or alignment artifact which provides less information about the real nucleotide distribution at this position. We set the percentage of N and - threshold to < 0.25 (indicated by the red vertical line in this plot) to keep the most informative group of sites in the genome).

The left hand side of the plot shows that there are over 100 positions with an entropy value around 1 (here we show the counts of entropy values greater than 0.1 because most of the positions in viral genomes have no variation so far and thus leads to 0 entropy at those positions—the peak at 0 masked entropy is not shown in full). However, most of those positions have high entropy because there are high percentage of N and - at those positions across all sequences in our dataset. The right hand side of the plot shows that there are a large amount of positions with high frequency of N and - (the peak at around 1.0). On the other hand, the peak at around 0 represents high quality positions with a fewer number of N and - present. According to this figure, we set the percentage of N and - threshold to 0.25 to keep the most informative group of sites in the genome.

### Supplementary file 2 — Sequence notation [40]

### Supplementary file 3 — ISM inflation by error correction

### Supplementary file 4 — Pairwise Linkage disequilibrium between high linkage sites

This table shows significantly linked pairs of sites and their the pairwise *r*^2^ value of Linkage disequilibrium [45].

### Supplementary file 5 — Visualizations of original ISMs

The figures in this document uses the same color codes for the original ISMs as the corresponding compressed ISMs. We can see from the figures that major genetic patterns are preserved in compressed ISM system.

### Supplementary file 6 — ISM abundance table of 20 countries/regions

The raw counts for all ISMs in each of 20 countries/regions, as well as the date each ISM was first found in a sequence in that country/region.

### Supplementary file 7 — ISM abundance table of 25 U.S. states

The raw counts for all ISMs in each of 25 U.S. states, as well as the date each ISM was first found in a sequence in that location.

### Supplementary file 8 — Geographical distribution of the dominant subtypes in Italy

This figure shows the relative abundance in other countries of the most abundant subtype in Italy CCCGCCAGGGA (left) and the second-most abundant subtype in Italy CCCGCCAGGGG (right).

Based on publicly available sequence data from Italy, we found that Italy had two particularly abundant ISMs, CCCGCCAGGGA and CCCGCCAGGGG, as can be seen in the pie chart in Figure 5. The third-most abundant subtype shown in the chart (CCCTCCAAGTG) corresponds to cases that were linked to original exposure from China, which is consistent with the ISM being in common with one found in Hong Kong. This supplementary figure shows the relative abundance (proportion of total sequences in that country/region) of each of these two dominant subtypes of Italy in other countries/regions. As the plot shows, the outbreak in other European countries have generally involved the same viral subtypes as those which are most abundant in Italy, as defined by ISM. Indeed, initial reports of cases in various other European countries in late February 2020 were linked to travelers from Italy [68]. The subtypes which are predominant in Italy are found, however, at lower yet notable abundance in countries including Japan, Canada and Australia.

Somewhat surprisingly, though the Italy subtypes were found in other U.S. states, only 88 out of the 1478 sequences from New York in the data set had the same ISM as the two dominant subtypes in Italy (see Supplementary file 7 — ISM abundance table of 25 U.S. states). This suggests that the outbreak in New York may not be linked to travel exposure directly from Italy, but rather from another location in Europe, with the important caveat that some potential subtypes may not have been detected there (due to relatively low number of sequences available from Italy). Indeed, the dominant subtype in New York (TCCGCCAGTGG) was detected in 86 sequence from Iceland and only one of them linked to travel exposure in Italy. However, 30 out of 86 cases linked to exposure in Austria, 6 linked to UK, 2 linked to Denmark, and 1 linked to Germany. This further suggests that it was unlikely that the incidence of the TCCGCCAGTGG subtype in New York is connected to Italy but rather than elsewhere in Europe, but limited sequence coverage in Italy prevents more definitive inference. However, one of the dominant subtypes in Italy, CCCGCCAGGGG, is not abundant in East Asian regions such as Mainland China and Japan, as indicated in this supplementary figure.

### Supplementary file 9 — Relative abundance (%) of ISMs in DNA sequences from Australia as sampled over time

Australia shows growing subtype diversity as its cases increase over time. Initially, Australia’s sequences were dominated by two subtypes that were also substantially abundant in Mainland China (CCCGCCAAGGG and CCTGCCAAGGG). Later, another subtype (CCCTCCAAGGG) starts to emerge. This subtype was less relatively abundant in Mainland China but more highly abundant in sequences from Hong Kong and Singapore (see Figure 5). Then, starting with sequences obtained on February 27, 2020, and subsequently, more subtypes are seen to emerge in Australia that were not found in other Asian countries but were found in Europe. This pattern suggests a hypothesis that Australia may have had multiple independent viral transmissions from Mainland China — or, as noted in the previous discussion, potentially through transmissions from Iran — followed by potentially independent importation of the virus from Europe and North America.

### Supplementary file 10 — Relative abundance (%) of ISMs in DNA sequences from Canada as sampled over time

This figure shows that the earliest viral sequences in Canada included mostly subtypes found in Mainland China, with the same pattern in which there was a second, later subtype in common with Mainland China, which was also found in travel exposure from Iran (CCCTCCAAGGG). And, like in Australia, in Canada these few initial viral sequences were followed by a diversification of subtypes that including many in common in Europe and the United States. In sum, Australia and Canada show patterns that might be expected for smaller populations in countries with diverse and extensive travel connections.

### Supplementary file 11 — Relative abundance (%) of ISMs in DNA sequences from Mainland China as sampled over time

This figure reflects Mainland China’s containment of SARS-CoV-2, as seen in the initial growth in viral genetic diversity, followed by a flattening as fewer new cases were found (and correspondingly fewer new viral samples were sequenced).

### Supplementary file 12 — Relative abundance (%) of ISMs in DNA sequences from the Netherlands as sampled over time

### Supplementary file 13 — Relative abundance (%) of ISMs in DNA sequences from Spain as sampled over time

This figure shows, in Spain, the CCCGCCAAGGG subtype was also found in an early sequence data but not thereafter. And, in Spain, a unique subtype has emerged that is not found in abundance in any other country.

### Supplementary file 14 — Temporal dynamics of individual viral subtypes across different regions

This figure shows that the reference genome subtype began to grow in abundance in Mainland China, before leveling off, and then being detected in the United States and Europe, and subsequently leveling off in those countries as well. In the case of Mainland China, that could be due to the substantial reduction in reported numbers of new infections and thus additional sequences being sampled. However, the other countries have continuing increases in reported infection as of the date of the data set, as well as substantially increasing numbers of sequences being sampled—making it less likely that the reference subtype (CCCGCCAAGGG) is simply being missed. In those cases, it appears from Figure 8, 9, and Supplementary file 12 — Relative abundance (%) of ISMs in DNA sequences from the Netherlands as sampled over time that in later times, other subtypes have emerged over time and are becoming increasingly abundant. One potential explanation is that the SARS-CoV-2 is an RNA virus and thus highly susceptible to mutation as transmissions occur [69]. Therefore, as transmissions have continued, the ISM associated with the reference sequence has been replaced by different ISMs due to these mutations. Another plausible explanation for such leveling off in a region is that the leveling off in relative abundance of the subtype represents containment of that subtype’s transmission while other subtypes continue to expand in that country or region. The latter could plausibly explain the pattern observed in the United States, where earlier subtypes connected to Asia did not increase in abundance while a putative endogenous subtype, as well as the dominant New York subtype, have significantly increased in abundance (see Figure 8 and accompanying discussion above). Further investigation and modeling of subtype distributions, as well as additional data, will be necessary to help resolve these questions — particularly in view of the caveats described below.

### Supplementary file 15 — Acknowledgements of sequences this research is based on

A list of sequences from GISAID’s EpiFlu Database on which this research is based and corresponding authors and laboratories.

## Footnotes

1 It is important to note that while we use the term “random” in the foregoing, in the biological context, a position may have a different base in different sequences due to selection pressure resulting in strains with different phenotypes, rather than purely random variation.

2 The latest report at the time of paper submission, run on June 22, 2020, with data up to June 17, 2020, can be found in https://github.com/EESI/ISM/blob/master/ISM-report-20200617-with_error_correction-compressed-SHORT-ISM.ipynb.

3 https://github.com/nextstrain/ncov

4 https://www.ncbi.nlm.nih.gov/

5 https://www.gisaid.org/

6 The sequences are of cDNA derived from viral RNA, so there is a T substituting for the U that would appear in the viral RNA sequence.

7 As discussed in the Results section, there are a few covarying positions that can be removed for a shorter ISM while still preserving the most of information. Therefore, for simplicity, we present SARS-CoV-2 subtypes in their short forms throughout this paper. In comparison with other methods, we include both the original ISM form and compressed ISM in the discussion. The visualizations of the original ISMs (including pie charts and time series charts) are available in Supplementary file 5 — Visualizations of original ISMs

## References

1. Hadfield J, Megill C, Bell SM, Huddleston J, Potter B, Callender C, et al. Nextstrain: real-time tracking of pathogen evolution. Bioinformatics. 2018 05;34(23):4121–4123. Available from: https://doi.org/10.1093/bioinformatics/bty407.

2. Li Q, Guan X, Wu P, Wang X, Zhou L, Tong Y, et al. Early Transmission Dynamics in Wuhan, China, of Novel Coronavirus–Infected Pneumonia. New England Journal of Medicine. 2020;382(13):1199–1207. Available from: https://doi.org/10.1056/NEJMoa2001316.

3. Tan W, Zhao X, Ma X, Wang W, Niu P, Xu W, et al. A novel coronavirus genome identified in a cluster of pneumonia cases—Wuhan, China 2019-2020. China CDC Weekly. 2020;2(4):61–2.

4. Benvenuto D, Giovanetti M, Salemi M, Prosperi M, Flora C, Alcantara L, et al. The global spread of 2019-nCoV: a molecular evolutionary analysis. Pathogens and Global Health. 2020 02;.

5. Sanjuán R, Nebot MR, Chirico N, Mansky LM, Belshaw R. Viral Mutation Rates. Journal of Virology. 2010;84(19):9733–9748. Available from: https://jvi.asm.org/content/84/19/9733.

6. Shu Y, McCauley J. GISAID: Global initiative on sharing all influenza data – from vision to reality. Eurosurveillance. 2017;22(13). Available from: https://www.eurosurveillance.org/content/10.2807/1560-7917.ES.2017.22.13.30494.

7. Forster P, Forster L, Renfrew C, Forster M. Phylogenetic network analysis of SARS-CoV-2 genomes. Proceedings of the National Academy of Sciences. 2020;117(17):9241–9243. Available from: https://www.pnas.org/content/117/17/9241.

8. Li X, Giorgi EE, Marichannegowda MH, Foley B, Xiao C, Kong XP, et al. Emergence of SARS-CoV-2 through recombination and strong purifying selection. Science Advances. 2020;6(27). Available from: https://advances.sciencemag.org/content/6/27/eabb9153.

9. Korber B, Fischer W, Gnanakaran S, Yoon H, Theiler J, Abfalterer W, et al. Spike mutation pipeline reveals the emergence of a more transmissible form of SARS-CoV-2. bioRxiv. 2020;Available from: https://www.biorxiv.org/content/early/2020/05/05/2020.04.29.069054.

10. Rambaut A, Holmes EC, Hill V, O’Toole Á, McCrone J, Ruis C, et al. A dynamic nomenclature proposal for SARS-CoV-2 to assist genomic epidemiology. bioRxiv. 2020;Available from: https://www.biorxiv.org/content/early/2020/04/19/2020.04.17.046086.

11. Zhao W, Song S, Chen M, Zou D, Ma L, Ma YK, et al. The 2019 novel coronavirus resource. Yi chuan = Hereditas. 2020;42 2:212–221.

12. Tang X, Wu C, Li X, Song Y, Yao X, Wu X, et al. On the origin and continuing evolution of SARS-CoV-2. National Science Review. 2020 03;Nwaa036. Available from: https://doi.org/10.1093/nsr/nwaa036.

13. Wang C, Liu Z, Chen Z, Huang X, Xu M, He T, et al. The establishment of reference sequence for SARS-CoV-2 and variation analysis. Journal of Medical Virology. 2020;n/a(n/a). Available from: https://onlinelibrary.wiley.com/doi/abs/10.1002/jmv.25762.

14. Sekizuka T, Itokawa K, Kageyama T, Saito S, Takayama I, Asanuma H, et al. Haplotype networks of SARS-CoV-2 infections in the Diamond Princess cruise ship outbreak. medRxiv. 2020;Available from: https://www.medrxiv.org/content/early/2020/03/27/2020.03.23.20041970.

15. Wang M, Li M, Ren R, Brave A, Werf Svd, Chen EQ, et al. International expansion of a novel SARS-CoV-2 mutant. medRxiv. 2020;Available from: https://www.medrxiv.org/content/early/2020/03/17/2020.03.15.20035204.

16. Jia Y, Yang C, Zhang M, Yang X, Li J, Liu J, et al. Characterization of eight novel full-length genomes of SARS-CoV-2 among imported COVID-19 cases from abroad in Yunnan, China. The Journal of Infection. 2020;.

17. Deng X, Gu W, Federman S, du Plessis L, Pybus OG, Faria NR, et al. Genomic surveillance reveals multiple introductions of SARS-CoV-2 into Northern California. Science (New York, Ny). 2020;.

18. Fauver JR, Petrone ME, Hodcroft EB, Shioda K, Ehrlich HY, Watts AG, et al. Coast-to-Coast Spread of SARS-CoV-2 during the Early Epidemic in the United States. Cell. 2020;181:990 – 996.e5.

19. Zhang L, Jackson CB, Mou H, Ojha A, Rangarajan ES, Izard T, et al. The D614G mutation in the SARS-CoV-2 spike protein reduces S1 shedding and increases infectivity. bioRxiv. 2020;Available from: https://www.biorxiv.org/content/early/2020/06/12/2020.06.12.148726.

20. Shen Z, Xiao Y, Kang L, Ma W, Shi L, Zhang L, et al. Genomic diversity of SARS-CoV-2 in Coronavirus Disease 2019 patients. Clinical Infectious Diseases. 2020 03;Ciaa203. Available from: https://doi.org/10.1093/cid/ciaa203.

21. Karamitros T, Papadopoulou G, Bousali M, Mexias A, Tsiodras S, Mentis A. SARS-CoV-2 exhibits intra-host genomic plasticity and low-frequency polymorphic quasispecies. bioRxiv. 2020;Available from: https://www.biorxiv.org/content/early/2020/03/28/2020.03.27.009480.

22. Grubaugh ND, Ladner JT, Lemey P, Pybus OG, Rambaut A, Holmes EC, et al. Tracking virus outbreaks in the twenty-first century. Nature Microbiology. 2019;4(1):10–19. Available from: https://doi.org/10.1038/s41564-018-0296-2.

23. Robinson ER, Walker TM, Pallen MJ. Genomics and outbreak investigation: from sequence to consequence. Genome Medicine. 2013;5(4):36. Available from: https://doi.org/10.1186/gm440.

24. Villabona-Arenas CJ, Hanage WP, Tully DC. Phylogenetic interpretation during outbreaks requires caution. Nature Microbiology. 2020;5(7):876–877. Available from: https://doi.org/10.1038/s41564-020-0738-5.

25. Year-letter Genetic Clade Naming for SARS-CoV-2 on Nextstain.org. Nextstrainorg. June 2, 2020;Available from: https://nextstrain.org/blog/2020-06-02-SARSCoV2-clade-naming.

26. Clarridge JE. Impact of 16S rRNA gene sequence analysis for identification of bacteria on clinical microbiology and infectious diseases. Clin Microbiol Rev. 2004;17:840–862.

27. Weisburg WG, Barns SM, Pelletier DA, Lane DJW. 16S ribosomal DNA amplification for phylogenetic study. Journal of bacteriology. 1991;173 2:697–703.

28. McDonald D, Birmingham A, Knight R. Context and the human microbiome. Microbiome. 2015 Nov;3(1):52. Available from: https://doi.org/10.1186/s40168-015-0117-2.

29. McDonald D, Hyde E, Debelius JW, Morton JT, Gonzalez A, Ackermann G, et al. American Gut: an Open Platform for Citizen Science Microbiome Research. mSystems. 2018;3(3). Available from: https://msystems.asm.org/content/3/3/e00031-18.

30. Gregory Caporaso J, Kuczynski J, Stombaugh J, Bittinger K, D Bushman F, K Costello E, et al. QIIME allows analysis of high-throughput community sequencing data. Nat Met 7: 335-336. Nature methods. 2010 04;7:335–6.

31. Cole JR, Wang Q, Fish JA, Chai B, McGarrell DM, Sun Y, et al. Ribosomal Database Project: data and tools for high throughput rRNA analysis. Nucleic Acids Research. 2013 11;42(D1):D633–D642. Available from: https://doi.org/10.1093/nar/gkt1244.

32. Eren AM, Maignien L, Sul WJ, Murphy LG, Grim SL, Morrison HG, et al. Oligotyping: differentiating between closely related microbial taxa using 16S rRNA gene data. Methods in Ecology and Evolution. 2013;4(12):1111–1119. Available from: https://besjournals.onlinelibrary.wiley.com/doi/abs/10.1111/2041-210X.12114.

33. Batista MVA, Ferreira TAE, Freitas AC, Balbino VQ. An entropy-based approach for the identification of phylogenetically informative genomic regions of Papillomavirus. Infection, Genetics and Evolution. 2011;11(8):2026 – 2033. Available from: http://www.sciencedirect.com/science/article/pii/S1567134811003236.

34. Dawy Z, Goebel B, Hagenauer J, Andreoli C, Meitinger T, Mueller JC. Gene mapping and marker clustering using Shannon’s mutual information. IEEE/ACM Transactions on Computational Biology and Bioinformatics. 2006;3:47–56.

35. Shannon CE. A mathematical theory of communication. The Bell System Technical Journal. 1948;27(3):379–423.

36. Crooks GE, Hon GC, Chandonia JM, Brenner SE. WebLogo: a sequence logo generator. Genome research. 2004;14 6:1188–90.

37. Bhowmik D, Pal S, Lahiri A, Talukdar A, Paul S. Emergence of multiple variants of SARS-CoV-2 with signature structural changes. bioRxiv. 2020;Available from: https://www.biorxiv.org/content/early/2020/04/29/2020.04.26.062471.

38. Katoh K, Standley DM. MAFFT Multiple Sequence Alignment Software Version 7: Improvements in Performance and Usability. Molecular Biology and Evolution. 2013 01;30(4):772–780. Available from: https://doi.org/10.1093/molbev/mst010.

39. Towns J, Cockerill T, Dahan M, Foster I, Gaither K, Grimshaw A, et al. XSEDE: Accelerating Scientific Discovery. Computing in Science Engineering. 2014 Sep;16(5):62–74.

40. Nomenclature for incompletely specified bases in nucleic acid sequences. Recommendations 1984. Nomenclature Committee of the International Union of Biochemistry (NC-IUB). Proceedings of the National Academy of Sciences. 1986;83(1):4–8. Available from: https://www.pnas.org/content/83/1/4.

41. Hunter JD. Matplotlib: A 2D graphics environment. Computing in Science & Engineering. 2007;9(3):90–95.

42. Inc PT. Collaborative data science. Montreal, QC: Plotly Technologies Inc.; 2015. Available from: https://plot.ly.

43. P Legendre LL. Numerical Ecology, Volume 24. Elsevier; 2008.

44. Novembre J, Johnson TB, Bryc K, Kutalik Z, Boyko AR, Auton A, et al. Genes mirror geography within Europe. Nature. 2008;456:98–101.

45. Beals EW. Bray-Curtis Ordination: An Effective Strategy for Analysis of Multivariate Ecological Data. In: Advances in Ecological Research. Elsevier; 1984. p. 1–55. Available from: https://doi.org/10.1016%2Fs0065-2504%2808%2960168-3.

46. Rosenberg A, Hirschberg J. V-Measure: A Conditional Entropy-Based External Cluster Evaluation Measure. In: EMNLP-CoNLL; 2007..

47. Korber B, Fischer WM, Gnanakaran S, Yoon H, Theiler J, Abfalterer W, et al. Tracking changes in SARS-CoV-2 Spike: evidence that D614G increases infectivity of the COVID-19 virus. Cell. XXXX 2020/07/02;Available from: https://doi.org/10.1016/j.cell.2020.06.043.

48. Grubaugh ND, Hanage WP, Rasmussen AL. Making sense of mutation: what D614G means for the COVID-19 pandemic remains unclear. Cell. XXXX 2020/07/02;Available from: https://doi.org/10.1016/j.cell.2020.06.040.

49. Kirchdoerfer RN, Ward AB. Structure of the SARS-CoV nsp12 polymerase bound to nsp7 and nsp8 co-factors. Nature Communications. 2019;10(1):2342. Available from: https://doi.org/10.1038/s41467-019-10280-3.

50. Ou X, Liu Y, Lei X, Li P, Mi D, Ren L, et al. Characterization of spike glycoprotein of SARS-CoV-2 on virus entry and its immune cross-reactivity with SARS-CoV. Nature Communications. 2020;11(1):1620. Available from: https://doi.org/10.1038/s41467-020-15562-9.

51. Walls AC, Park YJ, Tortorici MA, Wall A, McGuire AT, Veesler D. Structure, Function, and Antigenicity of the SARS-CoV-2 Spike Glycoprotein. Cell. XXXX 2020/04/06;Available from: https://doi.org/10.1016/j.cell.2020.02.058.

52. Yu IM, Gustafson CLT, Diao J, Burgner JWI, Li Z, qiang Zhang J, et al. Recombinant severe acute respiratory syndrome (SARS) coronavirus nucleocapsid protein forms a dimer through its C-terminal domain. The Journal of biological chemistry. 2005;280 24:23280–6.

53. To KKW, yin Tsang OT, shing Leung W, Tam AR, chiu Wu T, Lung DC, et al. Temporal profiles of viral load in posterior oropharyngeal saliva samples and serum antibody responses during infection by SARS-CoV-2: an observational cohort study. The Lancet Infectious diseases. 2020;.

54. Chan JFW, Kok KH, Zhu Z, Chu H, To KKW, Yuan S, et al. Genomic characterization of the 2019 novel human-pathogenic coronavirus isolated from a patient with atypical pneumonia after visiting Wuhan. Emerging Microbes & Infections. 2020;9(1):221–236. PMID: 31987001. Available from: https://doi.org/10.1080/22221751.2020.1719902.

55. Yuen KS, Ye ZW, Fung SY, Chan CP, Jin DY. SARS-CoV-2 and COVID-19: The most important research questions. Cell & Bioscience. 2020;10(1):40. Available from: https://doi.org/10.1186/s13578-020-00404-4.

56. Lewontin RC. On measures of gametic disequilibrium. Genetics. 1988;120 3:849–52.

57. Isabel S, Graña-Miraglia L, Gutierrez JM, Bundalovic-Torma C, Groves HE, Isabel MDR, et al. Evolutionary and structural analyses of SARS-CoV-2 D614G spike protein mutation now documented worldwide. bioRxiv. 2020;.

58. Bhattacharyya C, Das C, Ghosh A, Singh AK, Mukherjee S, Majumder PP, et al. Global Spread of SARS-CoV-2 Subtype with Spike Protein Mutation D614G is Shaped by Human Genomic Variations that Regulate Expression of TMPRSS2 and MX1 Genes. bioRxiv. 2020;.

59. Wu F, Zhao S, Yu B, mei Chen Y, Wang W, Song Z, et al. A new coronavirus associated with human respiratory disease in China. Nature. 2020;579:265 – 269.

60. Lauer SA, Grantz KH, Bi Q, Jones FK, Zheng Q, Meredith HR, et al. The Incubation Period of Coronavirus Disease 2019 (COVID-19) From Publicly Reported Confirmed Cases: Estimation and Application. Annals of Internal Medicine. 2020 May;172(9):577–582. Available from: https://doi.org/10.7326/M20-0504.

61. Lorenzo-Redondo R, Nam HH, Roberts SC, Simons LM, Jennings LJ, Qi C, et al. A Unique Clade of SARS-CoV-2 Viruses is Associated with Lower Viral Loads in Patient Upper Airways. medRxiv : the preprint server for health sciences. 2020;.

62. Gonzalez-Reiche AS, Hernandez MM, Sullivan MA, Ciferri B, Alshammary H, Obla A, et al. Introductions and early spread of SARS-CoV-2 in the New York City area. medRxiv. 2020;.

63. Yu WB, da Tang G, Zhang L, Corlett RT. Decoding the evolution and transmissions of the novel pneumonia coronavirus (SARS-CoV-2 / HCoV-19) using whole genomic data. Zoological Research. 2020;41:247 – 257.

64. Worobey M, Pekar JE, Larsen BB, Nelson MI, Hill V, Joy JB, et al. The emergence of SARS-CoV-2 in Europe and the US. bioRxiv. 2020;.

65. İlker Karacan, Akgun TK, Ağaoğlu NB, Irvem A, Alkurt G, Yildiz J, et al. The origin of SARS-CoV-2 in Istanbul: Sequencing findings from the epicenter of the pandemic in Turkey. Northern Clinics of Istanbul. 2020;7:203 – 209.

66. Miller D, Martin MA, Harel N, Kustin T, Tirosh O, Meir M, et al. Full genome viral sequences inform patterns of SARS-CoV-2 spread into and within Israel. medRxiv. 2020;.

67. Price MN, Dehal PS, Arkin AP. FastTree 2 – Approximately Maximum-Likelihood Trees for Large Alignments. PLoS ONE. 2010;5.

68. Coronavirus: Outbreak spreads in Europe from Italy, available at https://www.bbc.com/news/world-europe-51638095, last accessed 2020-04-05. BBC News. February 26, 2020;Available from: https://www.bbc.com/news/world-europe-51638095.

69. Moya A, Holmes EC, González-Candelas F. The population genetics and evolutionary epidemiology of RNA viruses. Nature Reviews Microbiology. 2004;2(4):279–288. Available from: https://doi.org/10.1038/nrmicro863.

