## supplemental file 5 for "Genetic Grouping of SARS-CoV-2 Coronavirus Sequences using Informative Subtype Markers for Pandemic Spread Visualization"

### Visualizations of original ISMs

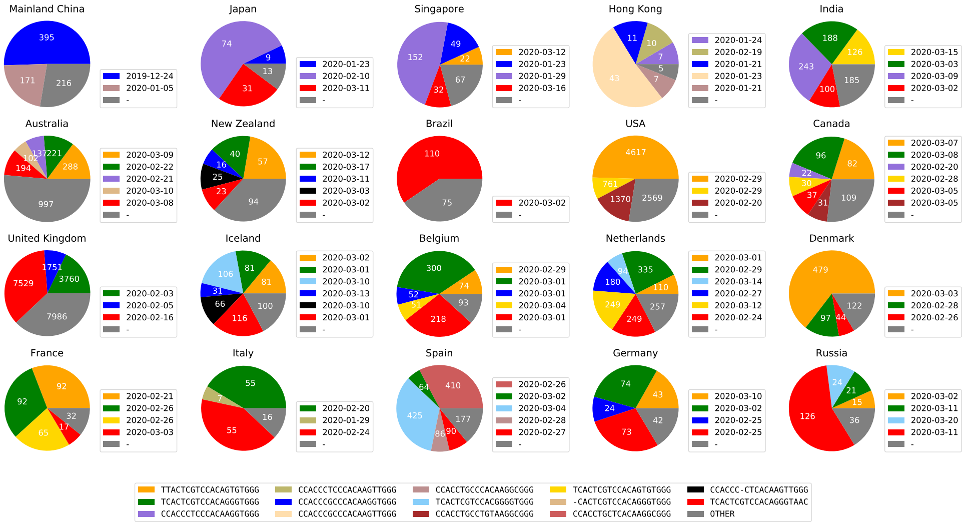


**Fig 1.** Major subtypes in countries/regions with the most sequences (indicating date subtype was first sequenced in that country/region). Subtypes with less than 5% abundance are plotted as "OTHER". The raw counts for all ISMs in each country/region, as well as the date each ISM was first found in a sequence in that country/region

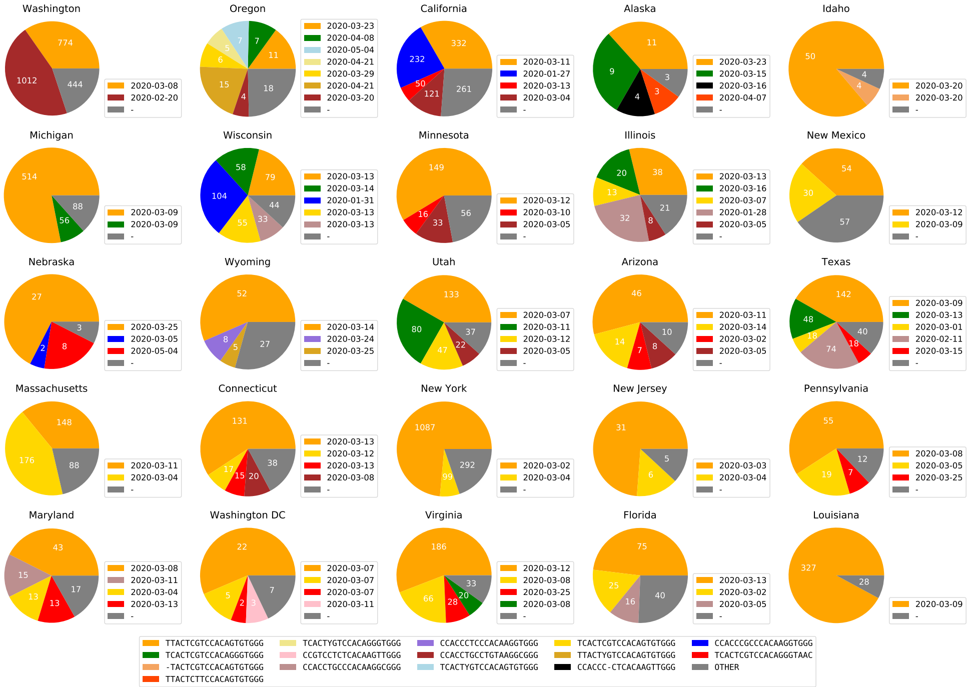


**Fig 2.** Viral subtype distribution in the United States, showing the 25 states with the most sequence submissions. Subtypes with less than 5% abundance are plotted as Other. The raw counts for all ISMs in each state, as well as the date each ISM was first found in a sequence in that state

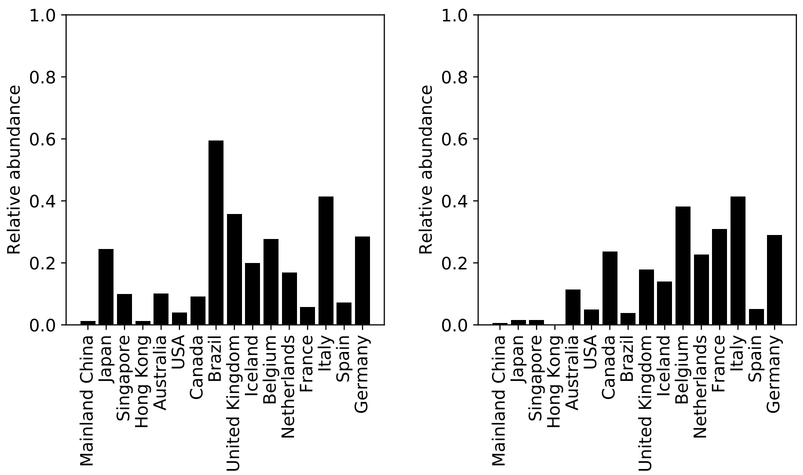


**Fig 3.** Relative abundance in other countries of the second-most abundant subtype in Italy, `TCACTCGTCCACAGGGTAAC` (left) and most abundant subtype in Italy `TCACTCGTCCACAGGGTGGG` (right).

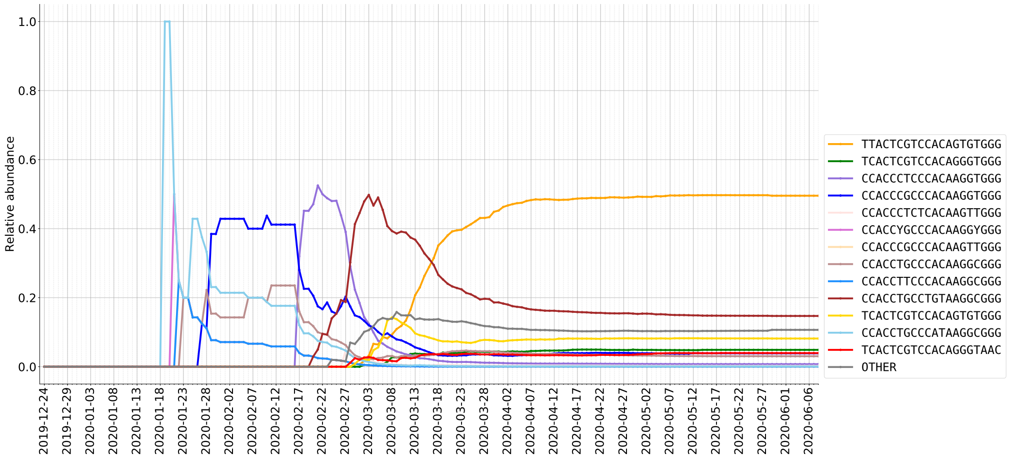


**Fig 4.** Relative abundance (%) of ISMs in DNA sequences from USA as sampled over time.

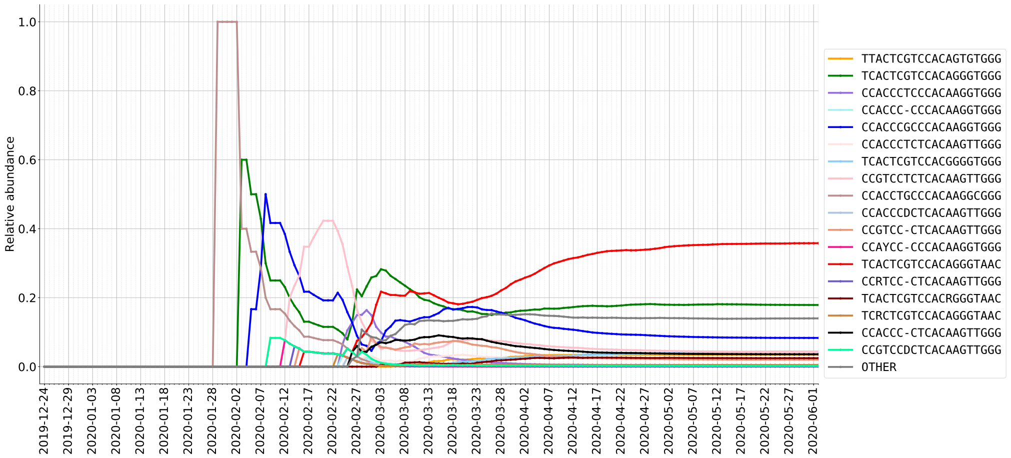


**Fig 5.** Relative abundance (%) of ISMs in DNA sequences from USA as sampled over time.

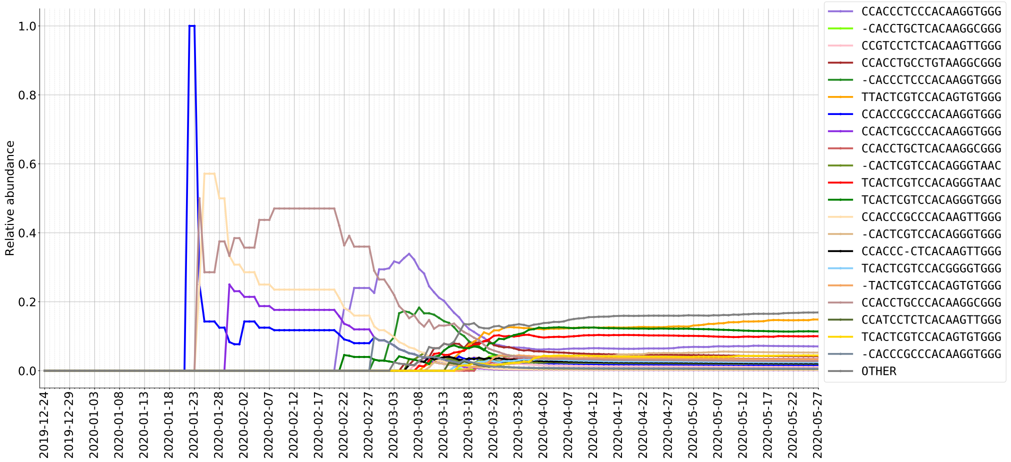


**Fig 6.** Relative abundance (%) of ISMs in DNA sequences from Australia as sampled over time.

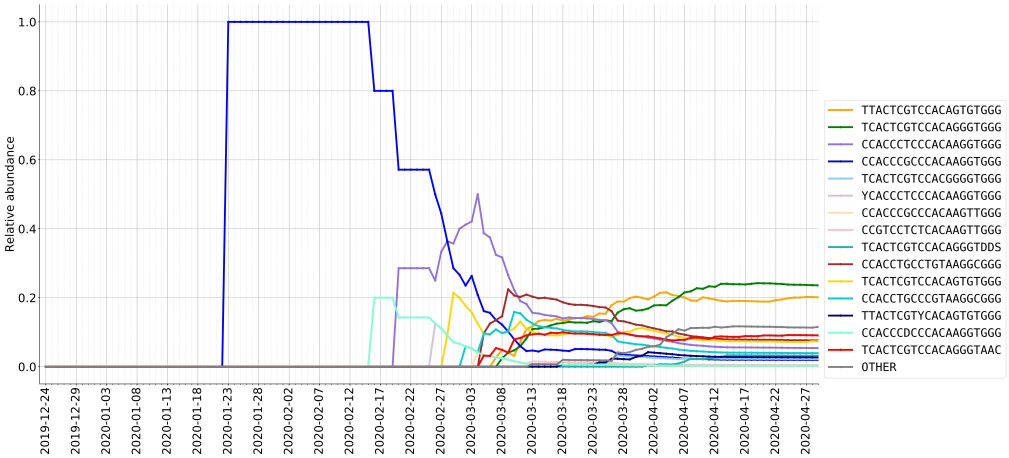


**Fig 7.** Relative abundance (%) of ISMs in DNA sequences from Canada as sampled over time.

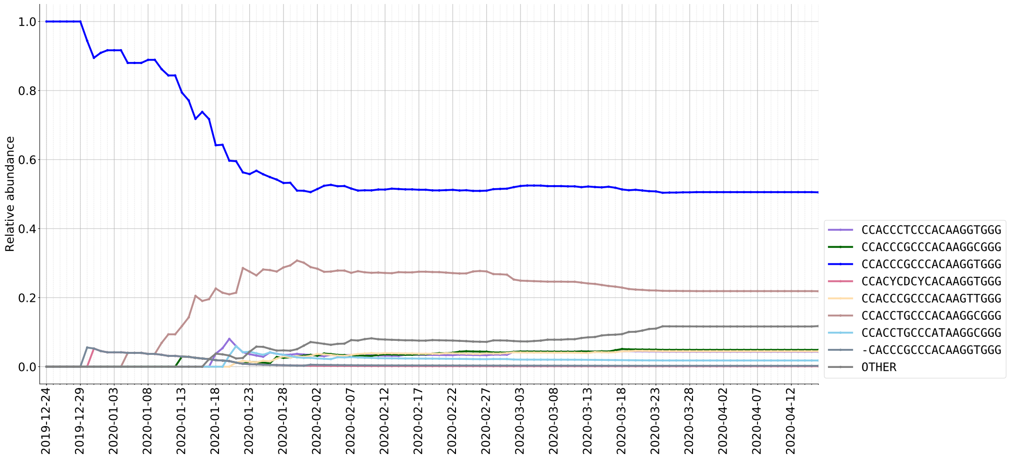


**Fig 8.** Relative abundance (%) of ISMs in DNA sequences from Mainland China as sampled over time.

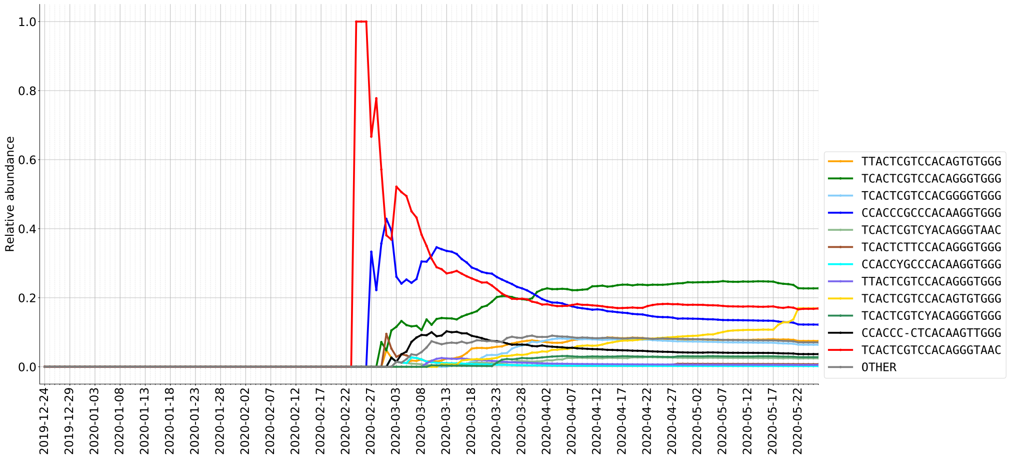


**Fig 9.** Relative abundance (%) of ISMs in DNA sequences from Netherlands as sampled over time.

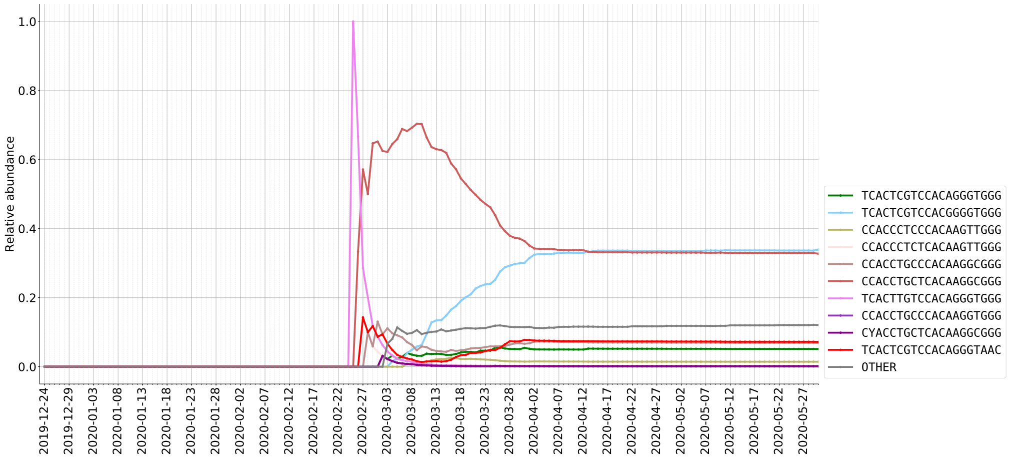


**Fig 10.** Relative abundance (%) of ISMs in DNA sequences from Spain as sampled over time.
