## Supplementary figures and images for "Genetic Grouping of SARS-CoV-2 Coronavirus Sequences using Informative Subtype Markers for Pandemic Spread Visualization"

### supplemental file 1

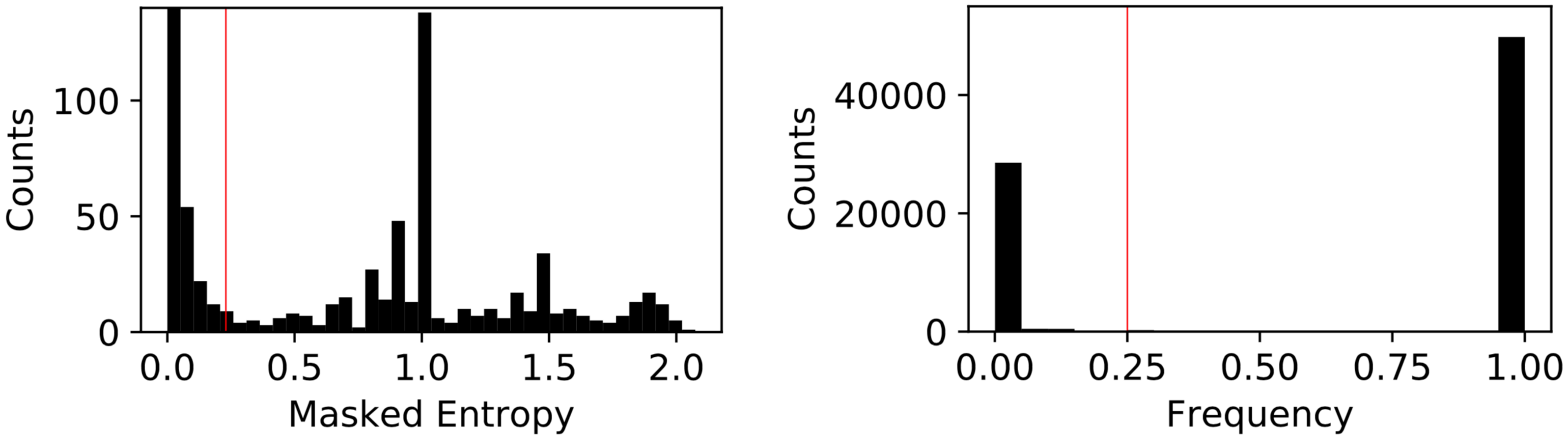

### supplemental file 8

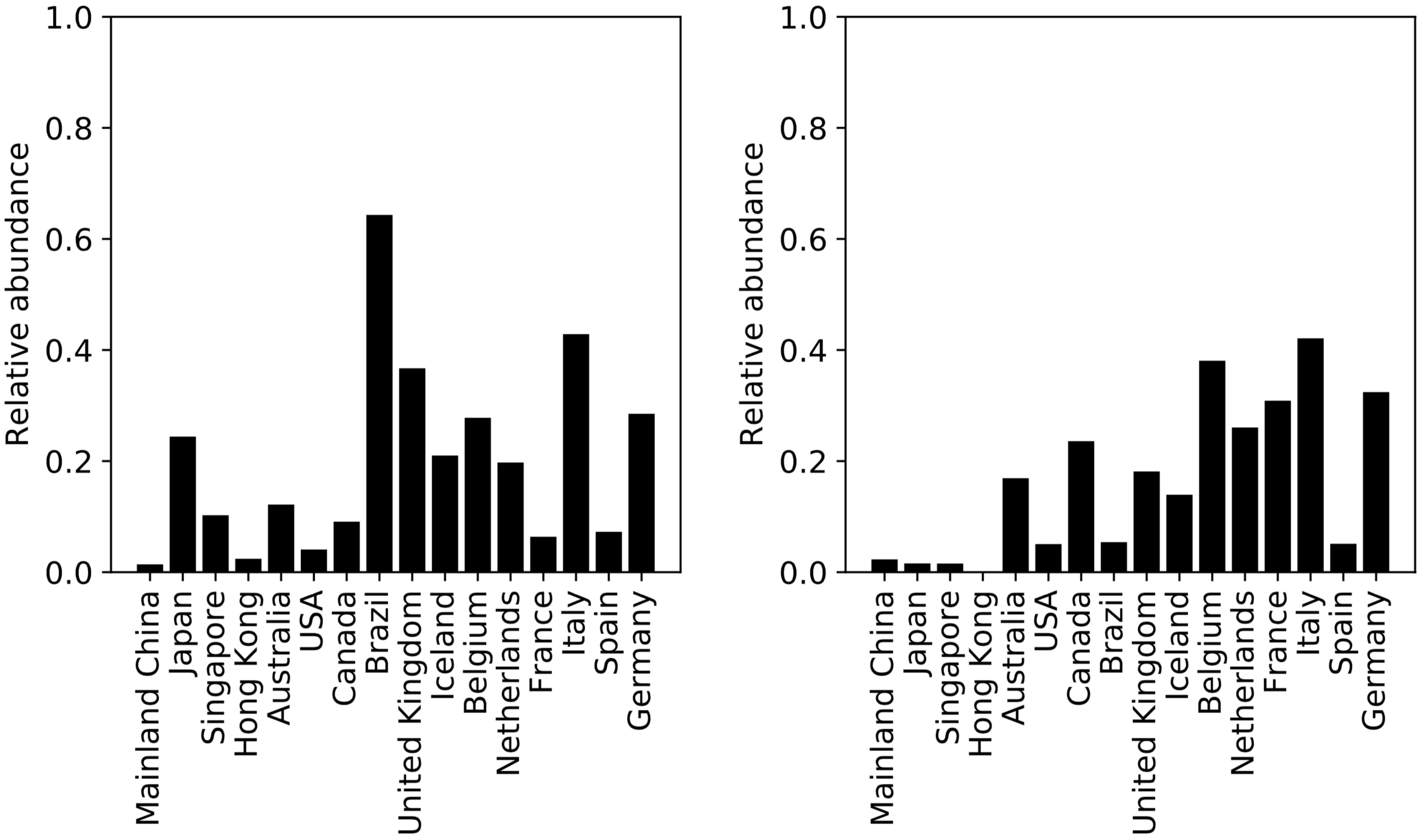

### supplemental file 9

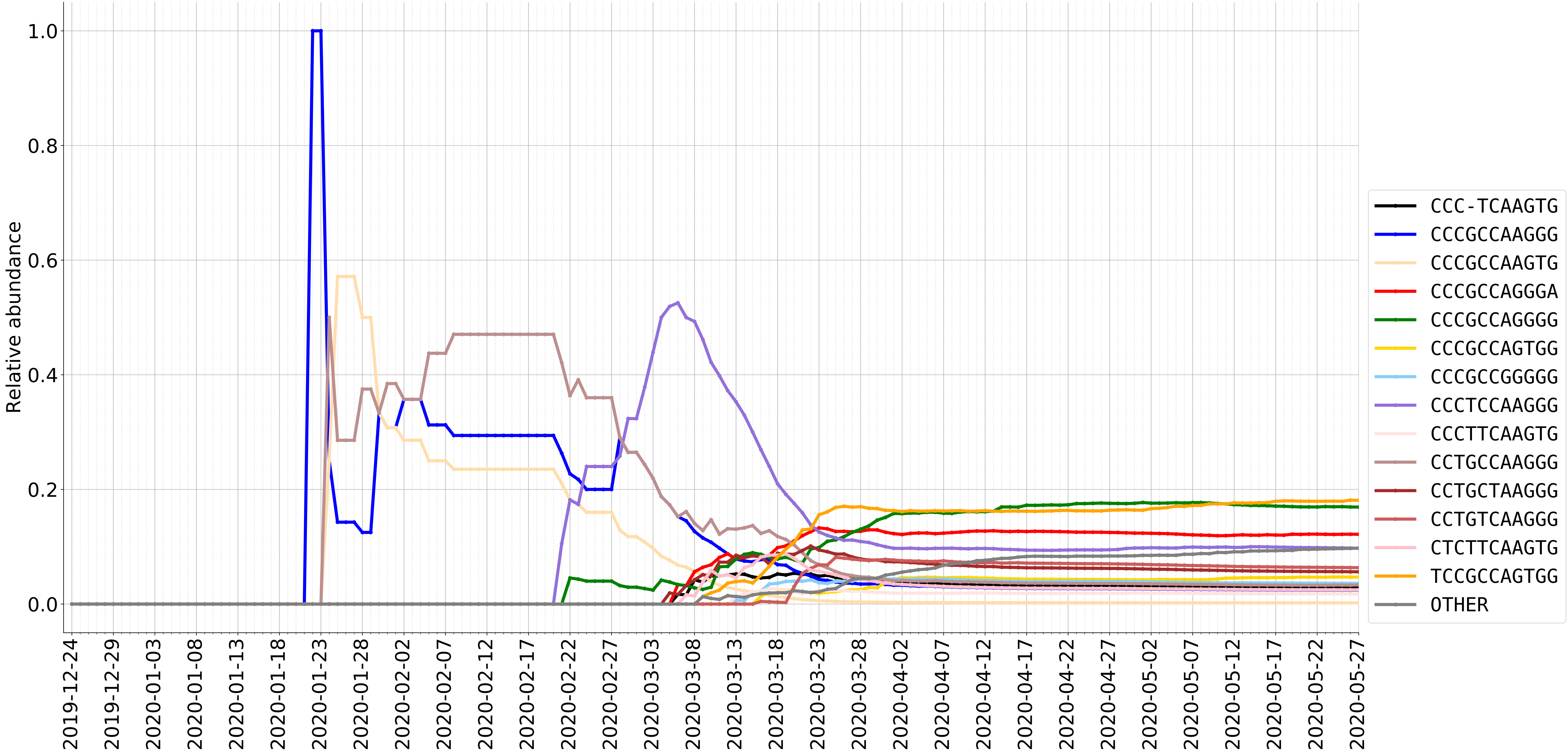

### supplemental file 10

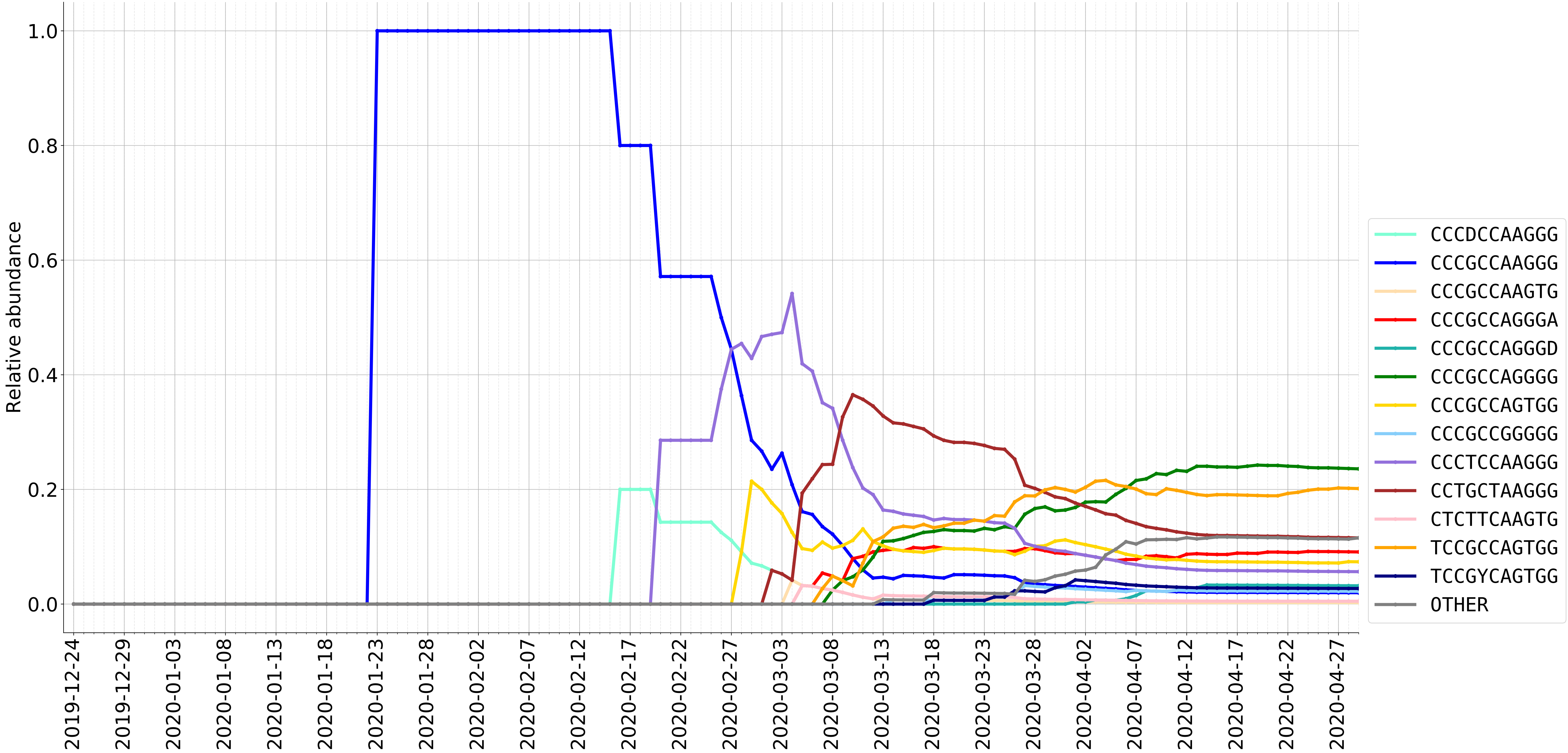

### supplemental file 11

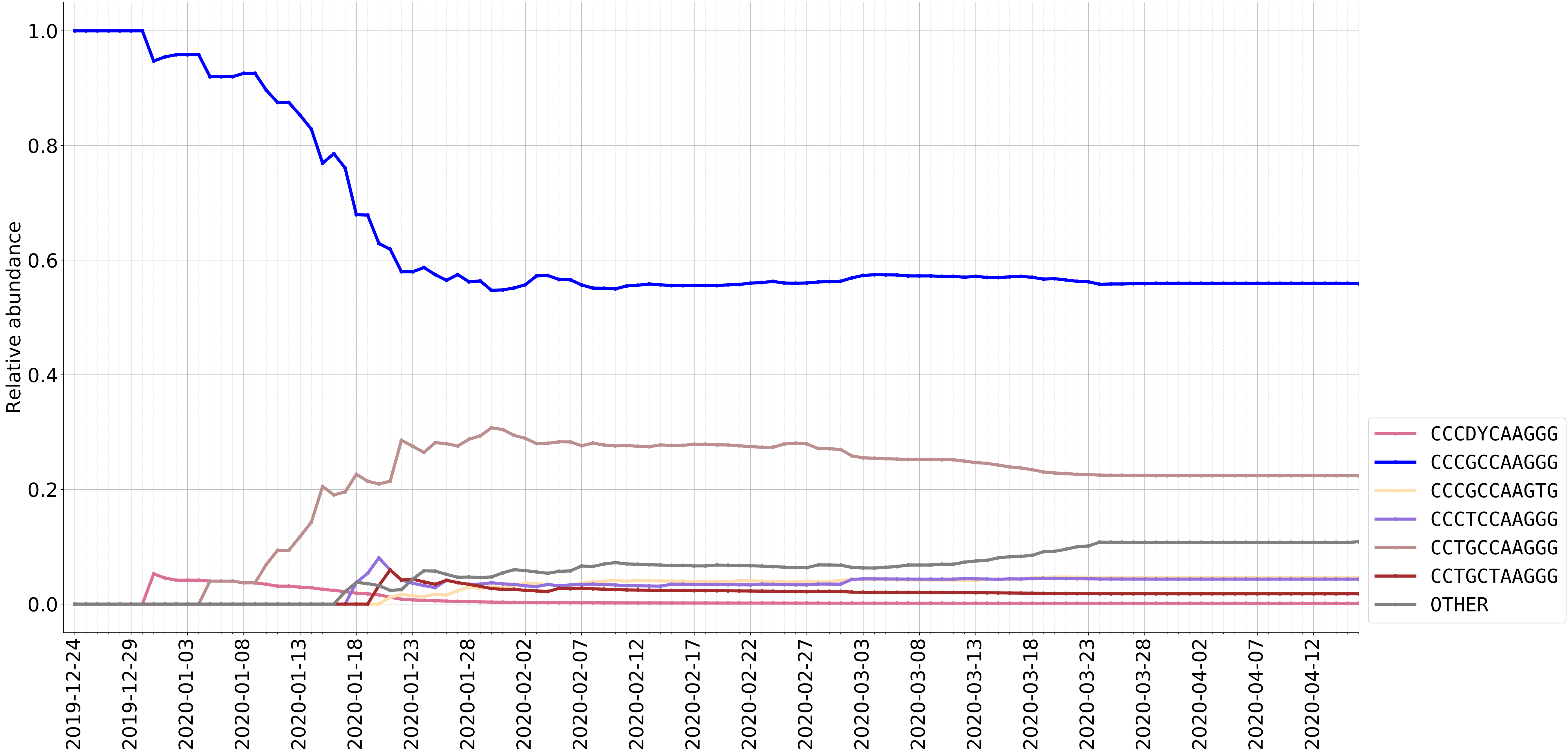

### supplemental file 12

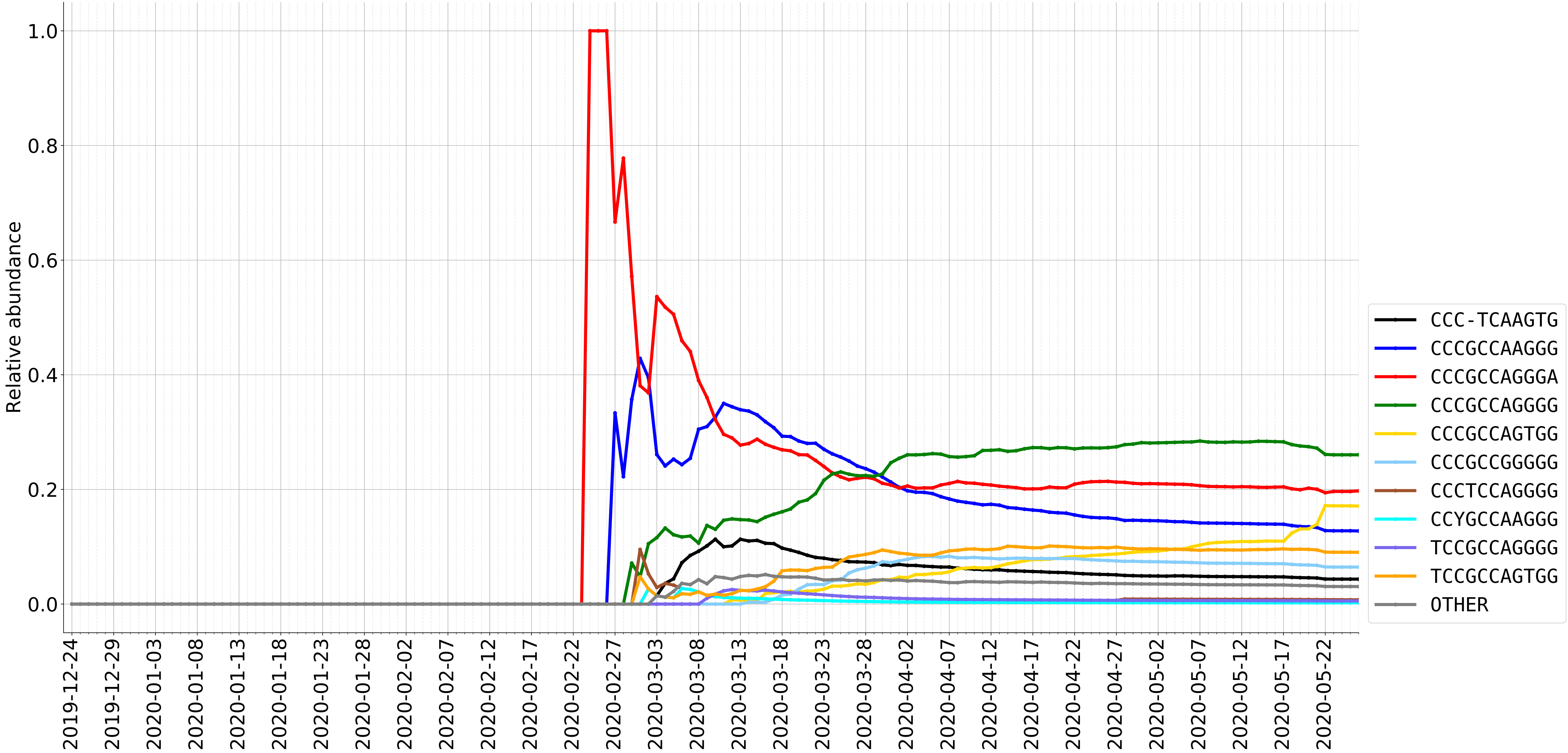

### supplemental file 13

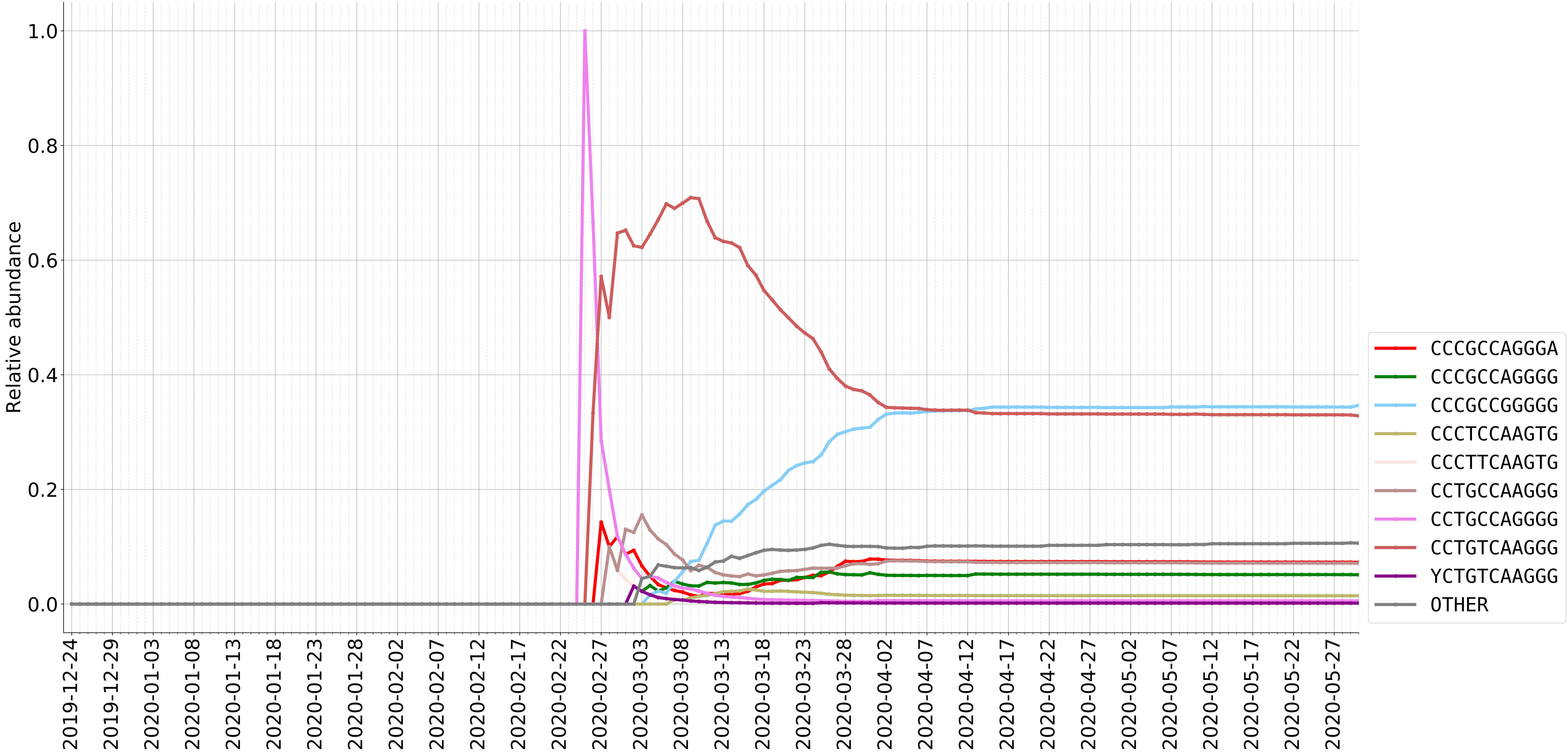

### supplemental file 14

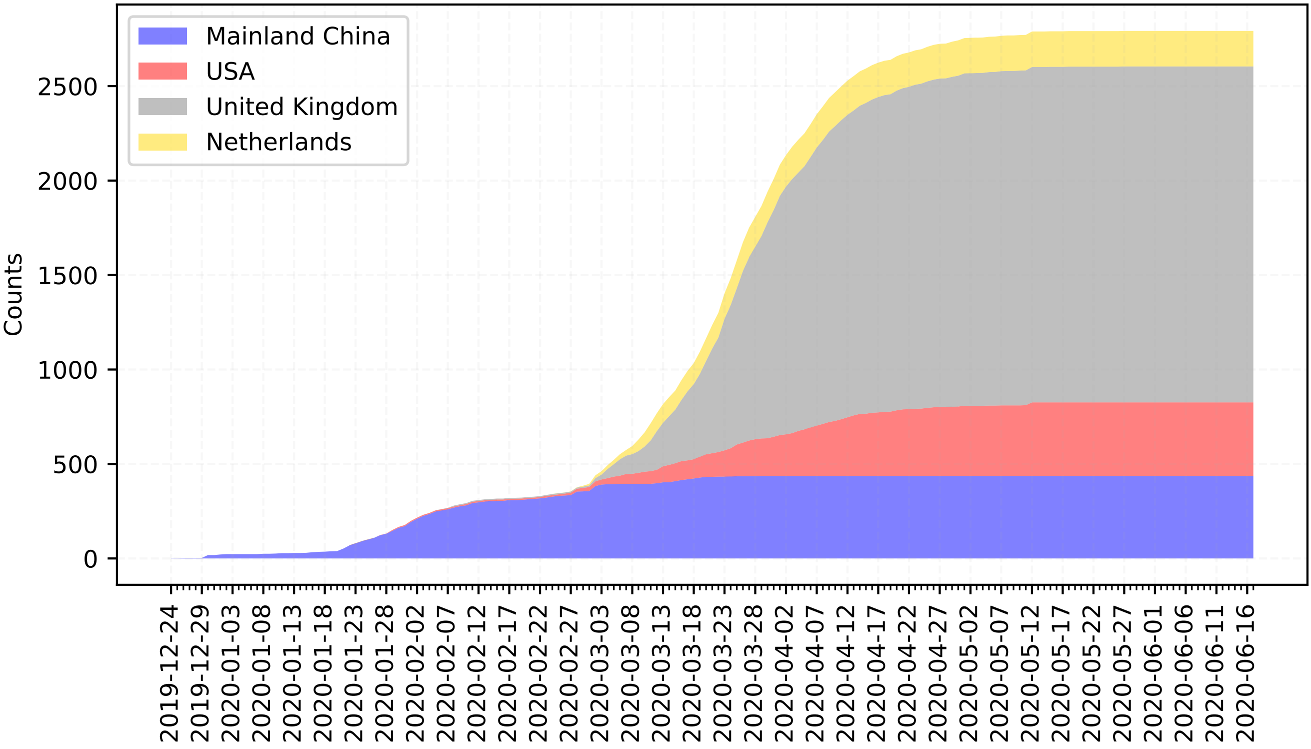
